# Highlighting strain rate dependent vibrational behavior of electrospun bundles for tendon/ligament and enthesis fascicle tissue regeneration

**DOI:** 10.64898/2026.01.14.699441

**Authors:** Alessandra Di Lorenzo, Tim Ten Brink, Gregorio Marchiori, Gianluca Giavaresi, Lorenzo Moroni, Martijn van Griensven, Alberto Sensini

## Abstract

Enthesis tissue engineering aims to develop scaffolds that replicate the mechanical and structural gradients of the tendon/ligament-bone interface. Among the different biofabrication techniques, electrospinning is surely one of the most promising to fabricate morpho-mechanically relevant enthesis fascicle-inspired scaffolds. An interesting and totally unexplored characteristic of these nanofibrous scaffolds is their ability, when mechanically tested, to produce/transmit strain rate and nanofiber fracture-dependent mechanical vibrations, which can potentially influence surrounding tissues and cells. This study develops a method to investigate how scaffold geometry and material affect vibrational behavior under mechanical stimulation. Electrospun bundles of poly(L-lactic) acid/collagen type I (PLLA/Coll) were fabricated to mimic the fibrocartilage, the enthesis junction, and the tendon/ligament regions, while block copolymer poly(ethylene oxide terephthalate)-poly(butylene terephthalate) (PEOT-PBT) bundles represented only the tendon/ligament. Scaffolds were morphologically and mechanically characterized, including strain rate-dependent vibrational response. Scanning electron microscopy confirmed distinct fiber architectures. Under monotonic tensile tests to failure, scaffolds exhibited strain rate-dependent mechanical behavior, with PLLA/Coll bundles showing dominant vibrational frequencies up to 4.2 ± 0.9 Hz with a scaffolds’ geometry-dependent manner. PEOT-PBT scaffolds instead, displayed higher vibration attenuation, with dominant frequencies peaking at 0.539 ± 0.063 Hz. They also showed lower tensile properties, reflecting a different mechanical and vibrational profile respect to PLLA/Coll bundles. Integrating vibrational characterization with mechanical testing offers a novel framework for designing scaffolds that more accurately reproduce the gradient mechanical environment of fibrous musculoskeletal tissues such as tendons/ligaments and their entheses. These findings highlight the potential of this combined approach to further increase the mechanical comprehension of electrospun scaffolds.

**Graphical Abstract:** 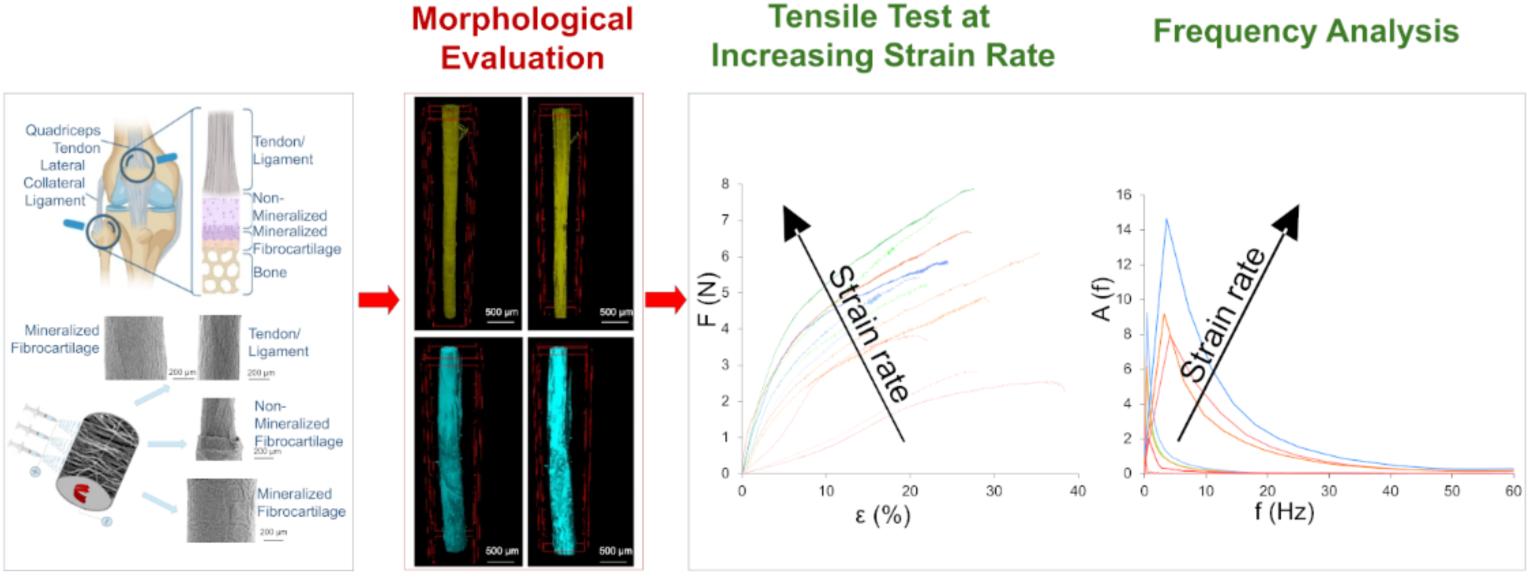

## 1. Introduction

Tendons and ligaments (T/L) are dense fibrous connective tissues essential for joint stability and musculoskeletal force transmission [1]. Their function relies on a hierarchical architecture of highly aligned collagen type I fibers, which confer tensile strength, viscoelasticity, and resistance to fatigue [1]. Their insertion site into bone, the enthesis, is a spatially graded interface composed of unmineralized and mineralized fibrocartilage that ensures efficient load transfer and mitigates stress concentrations [2,3]. This organized, anisotropic structure enables entheses to sustain high strain rates and complex multiaxial loads during movement, including impact and vibrational stimuli [4]. Regional variations in matrix composition and fiber alignment across these tissues (**Fig. 1A1, 1A2**) generate local differences in stiffness, damping capacity, and strain distribution, all of which are crucial to shock absorption, energy storage, and mechanical homeostasis [5,6]. During movement, T/L and their entheses experience not only tensile loads but also passive high-frequency vibrations generated during the daily life physiological activities and impacts [7–9]. These vibrations propagate along the extracellular matrix (ECM), transmitting mechanical cues to surrounding cells and thereby regulating strain distribution, cellular responses, and tissue remodelling [9,10]. Despite their functional specialization, T/L and entheses are frequently injured [11]. These injuries account for up to 50% of all sports-related traumas and are particularly prevalent in aging populations [11]. Healing is limited by the intrinsic avascularity and low cellularity of these tissues, which, together with their complex structural and mechanical environment, makes clinical repair particularly challenging [1].

**Fig. 1.**
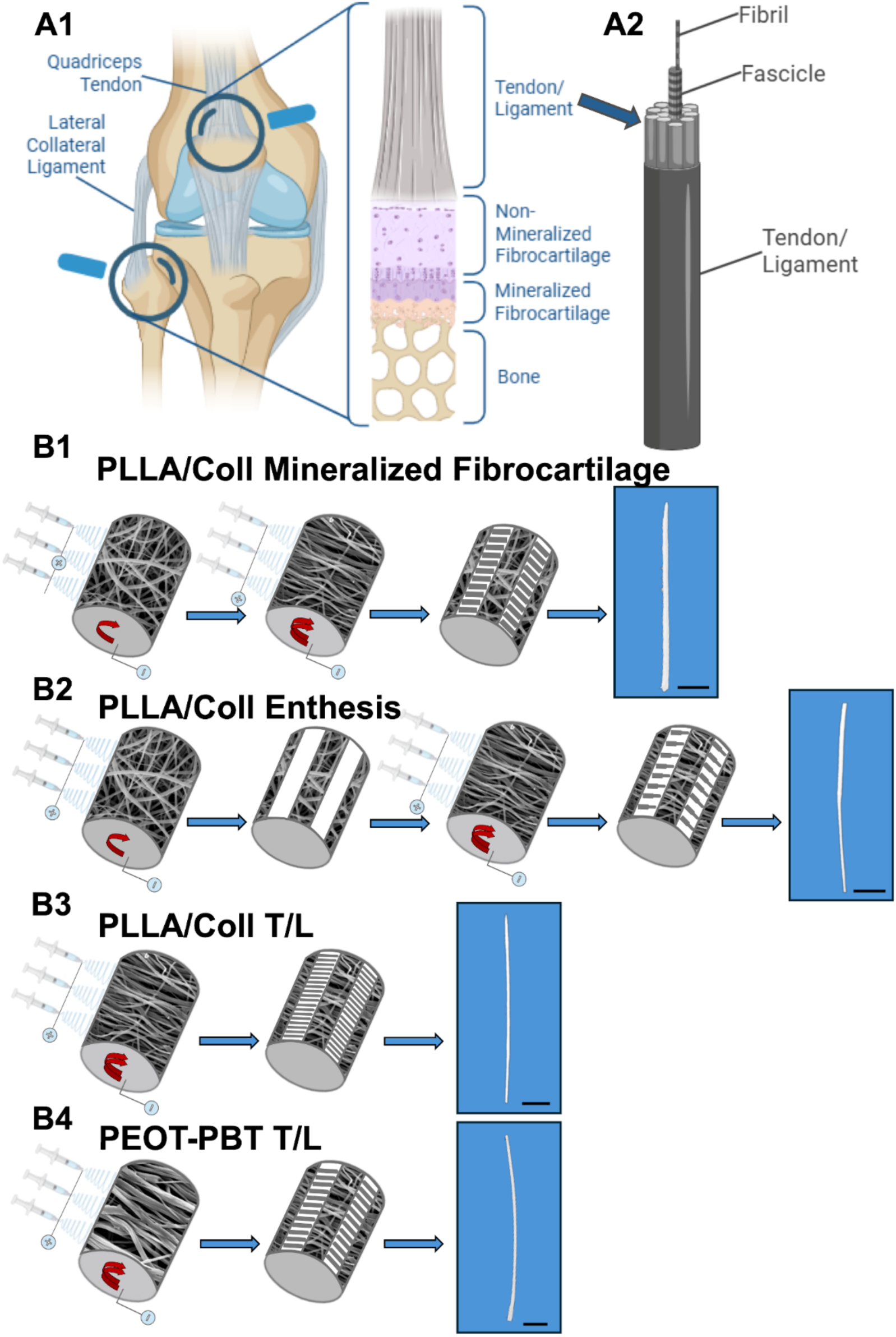
Structure of natural enthesis fascicles and biofabrication of electrospun bundles. A1) Example of natural enthesis and its structural composition; A2) hierarchical structure of T/L. Biofabrication steps for electrospun bundles’ production on a drum collector: B1) PLLA/Coll mineralized fibrocartilage-inspired bundles (FS); B2) PLLA/Coll enthesis-inspired bundles (ES); B3) PLLA/Coll T/L-inspired bundles (TLS); B4) PEOT-PBT T/L-inspired bundles (PTLS). All scale bars of typical bundles obtained are 5 mm long.

Conventional grafts and implants often fail to restore native functionality, resulting in poor biological integration and in mechanical mismatch [12,13]. Scaffolds that closely replicate the hierarchical fibrous architecture and mechanical properties of these tissues have been developed to address these shortcomings [14,15].

Electrospinning has emerged as a prominent biofabrication technique for T/L scaffolds due to its ability to generate micro- and nanofibrous structures, mimicking the collagen fibril-based ECM, essential for mechanical strength and cellular guidance [15]. By precisely tuning fiber diameter, orientation, and porosity, electrospun constructs can reproduce the structural complexity of the native ECM. The combination of electrospinning and careful material selection provides tuneable mechanical properties and promote favourable cell-scaffold interactions, supporting their potential for regenerative applications [16].

The viscoelastic properties of T/L electrospun constructs have been extensively characterized, but without focusing on their intrinsic vibrational behavior [8], [17,18]. Vibrational cues within engineered scaffolds have demonstrated to affect the cellular activity and ECM synthesis, underscoring the significance of these phenomena for regenerative outcomes [19,20]. The vibrational response of electrospun scaffolds to strain-rate dependent mechanical cues can be suitable for regenerative medicine [17,21]. Fast Fourier Transform (FFT) analysis provides a powerful tool to characterize the scaffolds frequency-dependent mechanical behavior [22–24]. It converts stress-strain measurements from the time to the frequency domain, revealing intrinsic features such as internal damping and energy dissipation. These characteristics depend on both construct architecture and composition, offering guidance for designing scaffolds that aim of replicating the vibrational environment of native tissues.

Several materials were investigated for T/L applications, including the block copolymer poly(ethylene oxide terephthalate)-poly(butylene terephthalate) (PEOT-PBT), composed of alternating hydrophilic and semi-aromatic segments [25,26–28]. This provided graded mechanical properties for osteotendinous tissue regeneration [27]. Blends of collagen type I (Coll) and poly(L-lactic acid) (PLLA) have also been explored to mimic both the structural and biochemical features of T/L and enthesis fascicles [29–32]. PLLA is a biodegradable aliphatic polyester with slow degradation kinetics that enhances the mechanical properties of collagen [33]. PLLA/collagen (PLLA/Coll) electrospun bundles in hierarchical-fiber configuration have demonstrated suitable mechanical performance for T/L land enthesis tissue engineering and the ability to promote cell alignment and ECM deposition [34]. PEOT-PBT and PLLA/Coll electrospun scaffolds have been characterized mechanically, but their vibrational response during progressive nanofiber rupture under tensile loading remains unexplored [27,30,31].

In this work, we developed electrospun scaffolds in a dual configuration: PLLA/Coll bundles replicating the non-mineralized fibrocartilage, the enthesis, and the T/L, and PEOT-PBT bundles tailoring exclusively to the T/L. This to investigate how the different materials’ stiffness can affect the vibrational components of scaffolds. We investigated their morphological biomimicry via scanning electron microscopy (SEM) and x-ray micro-computed tomography (microCT). Then, uniaxial tensile tests up to failure, at different strain rates, coupled with frequency-domain analysis were carried out, aiming to study the structure strain rate-dependent vibrational responses of the fibrous scaffolds.

## 2. Materials and Methods

### 2.1. Materials

The electrospun PLLA/Coll bundles were fabricated using acid-soluble collagen type I, extracted from bovine dermis (Kensey Nash Corporation DSM Biomedical, Exton, USA), and medical-grade poly(L-lactic) acid (PLLA; Purasorb PL18, Mw = 1.7-1.9 × 10⁵ g mol⁻¹; Corbion, Amsterdam, The Netherlands). The two polymers were blended in a 75:25 weight ratio and dissolved at a total concentration of 15% (w/v) in 1,1,1,3,3,3-hexafluoro-2-propanol (HFIP; Sigma-Aldrich, Saint Louis, USA).

The electrospun PEOT-PBT bundles were manufactured using a 200 mg·mL^-1^ PEOT-PBT copolymer formulation, prepared with a solvent blend of 75 % (v/v) of chloroform and 25 % (v/v) of 1,1,1,3,3,3-hexafluoro-2-propanol. This solution was kept at room temperature for 12 hours under agitation to ensure proper mixing.

### 2.2. Bundles preparation

Electrospinning was performed using commercial Fluidnatek systems (LE-100 and LE-500, Bioinicia, Spain) equipped with multi-needle assemblies (three needles per setup). For each setup, the polymer solution was loaded into three 5 mL plastic syringes (BD Biosciences, New Jersey, US) connected via PTFE tubing to stainless steel needles of 0.51 mm and 0.8 mm internal diameters for PLLA/Coll and PEOT-PBT solutions, respectively. Fibers were collected on a rotating aluminium drum (diameter = 200 mm, length = 300 mm), covered with polyethylene-coated paper (Turconi S.p.A, Ceriano Laghetto, Italy) to facilitate subsequent mat removal. Environmental conditions settled at a temperature of 25°C and a relative humidity of 25%.

The fabrication of nanofibrous bundles was inspired by the native hierarchical architecture and dimensional features of T/L fascicles and their enthesis. A composite design was implemented to recapitulate the transition from mineralized fibrocartilage to T/L through a conical junction region [35].

For PLLA/Coll scaffolds, electrospinning parameters were set to an applied voltage of 24 kV (needle) and -0.5 kV (collector), feed rate of 600 μL·h^−1^, and needle-collector distance of 200 mm. Fiber deposition was modulated by alternating drum rotation speeds. Three different bundles’ configurations were produced resembling the mineralized fibrocartilage fascicle region (FS), the T/L fascicle region (TLS) and the whole enthesis fascicle (ES) including both FS and TLS connected by a conical junction of aligned nanofibers (non-mineralized fibrocartilage inspired). Specifically, to produce the bundles, different biofabrication steps were adopted (using the electrospinning parameters previously showed):

- FS: PLLA/Coll nanofibers were spun for three hours on the drum rotating at low speed (50 rpm, peripheral speed = 0.52 m s^-1^), obtaining a mat of randomly arranged nanofibers, followed by additional three hours at high speed (2000 rpm; peripheral speed = 20.9 m s^-1^), obtaining, on top of the previous, a mat of circumferentially aligned nanofibers (**Fig. 1B1**). Mats were cut into strips (30 mm wide and 40 mm along the circumference) and manually wrapped along the drum axis obtaining bundles 40 mm long.

- ES: PLLA/Coll nanofibers were spun for three hours on the drum rotating at low speed (as for FS). The mat was divided in axial stripes (20 mm along the circumference) and alternatively removed from the drum. Then, three hours of electrospinning at high speed (as for SF) were carried out obtaining alternate stripes of FS and TLS regiones (of 20 mm along the circumference each). Stipes were finally cut (30 mm wide and 40 mm along the circumference) and manually wrapped along the drum axis obtaining bundles 40 mm long with the two regions connected by a conical non-mineralized fibrocartilage-inspired junction of aligned nanofibers (Fig.1B2).

- TLS: PLLA/Coll nanofibers were spun for three hours on the drum rotating at high speed (as for FS). The mat was cut in stripes (30 mm wide and 40 mm along the circumference) and manually wrapped along the drum axis obtaining bundles of aligned nanofibers 40 mm long (Fig. 1B3).

Finally, similarly as TLS, PEOT-PBT fiber mats were produced using a Fluidnatek LE-500 unit, applying an identical voltage scheme (24 kV needle, -0.5 kV collector), feed rate of 600 μL h^-1^, and a shorter needle-to-collector distance (120 mm). The needle assembly executed a linear scanning movement parallel to the drum, spanning 190 mm at 3600 mm min^-1^, ensuring uniform fiber deposition. The drum rotated at 2000 rpm to induce fiber alignment (Fig. 1B4). Post-electrospinning, mats were sectioned into strips (30 mm wide and 40 mm along the circumference) and manually wrapped along the collector axis to form bundles. These bundles simulated the T/L region (these specific bundles called PTLS) and were used as comparison with the previous ones. This integrated electrospinning approach enabled the fabrication of fibrous biomimetic bundles with controlled architecture, fiber orientation, and dimensions closely matching native tissue features.

### 2.3. Collagen crosslinking

Bundles of PLLA/Coll were chemically crosslinked to fix the collagen in nanofibers. Crosslinking was performed using N-(3-dimethylaminopropyl)-N′-ethylcarbodiimide hydrochloride (EDC), N-hydroxysuccinimide (NHS), and 95% ethanol (EtOH), all used as received (Sigma-Aldrich, Saint Louis, USA), for 24 hours at room temperature, following an established protocol [31]. After crosslinking, the samples were incubated in 0.1 M phosphate-buffered saline (PBS, pH 7.4) for 30 minutes, rinsed in distilled water over a 2-hour period with water replacement every 15 minutes, and subsequently air-dried under a chemical hood at room temperature for 24 hours [31].

### 2.4. Morphological investigation

To investigate the surface morphology of the bundles, SEM (JSM-IT200 InTouchScope™ JEOL, Japan) was employed using a voltage of 10 KeV. Prior to imaging, the specimens were gold sputtered to ensure conductivity, and observations were performed at an accelerating voltage of 10 kV. To assess the nanofiber orientation, SEM images were analysed using the Directionality plugin of the open-source ImageJ software.

To quantify the orientation of nanofibers, a previously validated procedure was adopted [36]. Specifically, the Local Gradient Orientation method, as described by Sensini *et al*. [36], was implemented. This method computes local gradients across grayscale SEM images to determine the predominant orientation of fibers. For each scaffold category, five SEM micrographs at 5000× magnification were analysed to generate orientation histograms, thereby quantifying the distribution of fibers within specified angular intervals.

To evaluate the three-dimensional morphology of the scaffolds, microCT (Skyscan 1172, Bruker, Belgium) was employed. Two simple bundles (PEOT-PBT) and two entheses (PLLA/Coll) were scanned with an applied voltage of 40 kV and current of 75 µA (no metal filter). The scan orbit was 360° with a rotation step of 0.2° and 4 frames averaged for each rotation angle, the voxel size was 9 µm, resulting in a scan duration of ⁓ 2 hours. Scan (i.e. projection) images were reconstructed in transversal images with a modified Feldkamp algorithm using the SkyScan Nrecon software accelerated by GPU, applying a level 5 ring artefact correction, without beam hardening correction and smoothing, by following consolidated protocols [30,37]. Each imaged object was divided into volumes of interest (VOI, height 3 mm) along the longitudinal direction: for bundle and for fibrocartilage or T/L regions, at different height levels; for enthesis, below/on/above junction (**Fig.2**). Three-dimensional fibrous arrangement was qualitatively highlighted using Skyscan CT-Vox rendering software, while porosity and diameter were calculated by Skyscan CT-Analyser software after binarization, for each VOI.

**Fig. 2.**
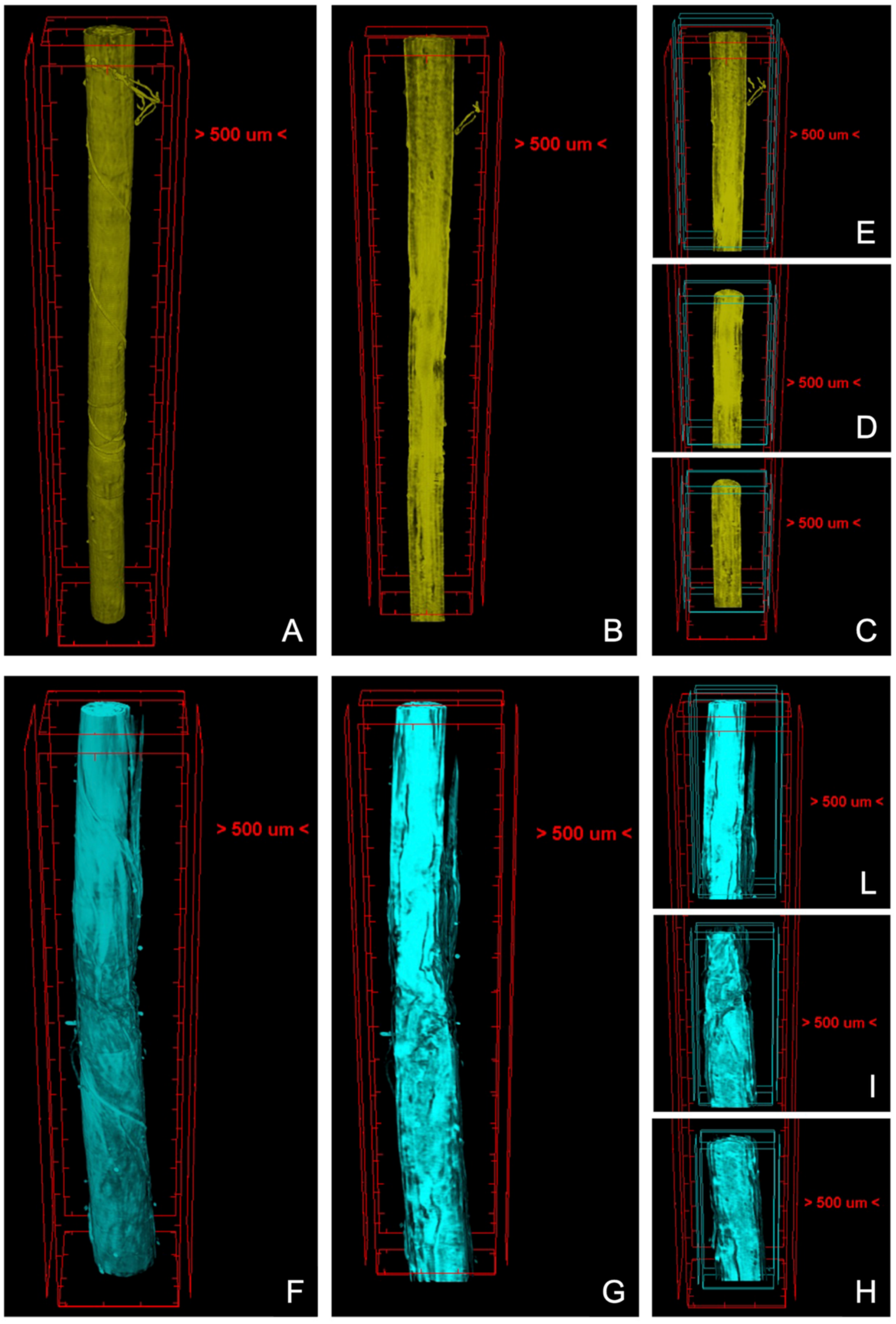
MicroCT image renderings (CT-Vox software, Skyscan) of: PEOT-PBT bundle as a whole (A), as longitudinally cut (B) and divided in bottom (C), middle (D), top (E) VOI; PLLA/Coll junction as a whole (F), as longitudinally cut (G), and divided in bottom (H), middle (I), top (L) VOI.

### 2.5. Mechanical investigation

The mechanical performances of scaffolds were assessed through uniaxial tensile tests conducted at room temperature using a mechanical testing machine (ElectroForce 3200, TA Instruments, New Castle, DE, USA) equipped with a ±44 N load cell. Five specimens per experimental group, including PTLS samples, were tested. Prior to testing, all samples were hydrated by immersion in phosphate-buffered saline (PBS, Sigma-Aldrich) for 2 minutes to simulate physiological moisture conditions.

Tensile tests were performed under displacement control following a monotonic ramp-to-failure protocol in accordance with ASTM D1414. Crosshead displacement was regulated using the WinTest 7 software (ElectroForce Systems Group, TA Instruments) to achieve predefined strain rates of 0.4% s⁻¹, 10% s⁻¹, and 100% s⁻¹ (bundles gauge length = 20 mm) [38]. These rates were selected to simulate physiological conditions ranging from quasi-static muscle loading (0.4% s⁻¹), to moderate dynamic activity such as fast walking or a slow run (10% s⁻¹), and high-rate events that may lead to T/L injury (100% s⁻¹), as previously reported [38–40]. Sampling rates were selected to ensure adequate temporal resolution at each strain rate: 20 Hz for 0.4% s⁻¹, 1000 Hz for 10% s⁻¹, and 2500 Hz for 100% s⁻¹.

Before the test, care was taken to secure crosshead stability and the elimination of any environmental vibrations or noise that may affect the mechanical test.

Mechanical properties were determined considering scaffold porosity, which was calculated from sample mass, geometry, and polymer density following established methods [41,42]. Briefly, the volume fraction was calculated as:

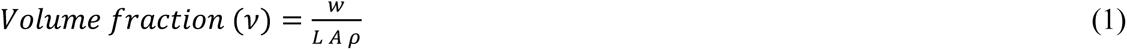

where w is the weight, *L* is the length, *A* is the cross-sectional area, and *ρ* the composite density of the specimen. The density (*ρ*) of the PLLA/Coll bundles was calculated according to the composite density formula [70]:

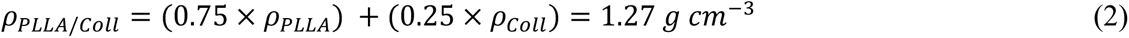

where *ρ_PLLA_* = 1.25 *g cm*^−3^ and *ρ_Coll_* = 1.34 *g cm*^−3^. For the PEOT-PBT instead the density of the bulk polymer was adopted (*ρ_PEOT_*_–*PBT*_ = 1.20 *g cm*^−3^). The percentage of porosity for each sample category (mean and standard deviation of n = 5 specimens per category) was assessed considering the following equation [41,42]:

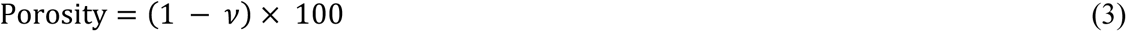

This expression accounts for the void fraction within the fibrous matrix, enabling a more accurate estimation of the material’s intrinsic mechanical response (i.e., net mechanical properties).

Force-displacement data were post-processed using a custom MATLAB routine to extract key mechanical parameters, including force at yield (F_Y_), yield strain (ε_Y_), yield stress (σ_Y_), stiffness (K), elastic modulus (E), force at failure (F_F_), failure strain (ε_F_), failure stress (σ_F_), work to yield (W_Y_), and work to failure (W_F_).

The low-strain region of the stress-strain curves was analysed to evaluate the mechanical response during the early loading regime. The inflection point (IF), defined as the transition from strain-stiffening to strain-softening behavior of the stress-strain curves, was identified using a previously validated MATLAB algorithm [39,40]. This point corresponds to the maximum curvature of the stress-strain curve and was determined as the zero of its second derivative [44]. From it, the following parameters were extracted: IF force (IFF), IF strain (IFε), IF stress apparent (IFσ_App_), and IF stress net (IFσ_Net_).

### 2.6 Frequency-domain vibration analysis

Mechanical vibration characteristics in scaffolds, induced by the imposed displacement, were quantified using a custom MATLAB routine that processed the force-time-series collected during tensile testing. The script applies a Fast Fourier Transform (FFT) to the time-domain stress and strain signals to extract frequency components, using the strain rate and sampling rate as input parameters. From the resulting frequency spectrum, 4 key parameters were computed: the dominant frequency, the signal amplitude at the dominant frequency, the wavelength, and the longitudinal wave speed.

The dominant frequency was defined as the frequency corresponding to the maximum amplitude in the FFT spectrum, representing the most energetically significant oscillation during mechanical wave propagation. The amplitude at the dominant frequency reflects the magnitude of the signal at this peak frequency and serves as an indicator of wave attenuation and energy dissipation within the material.

The wavelength of the dominant wave was estimated by analysing the periodicity of the signal in the time domain. Once both the dominant frequency and the wavelength were identified, the longitudinal wave speed was calculated using the relation:

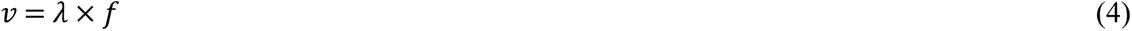

where *v* (m s^-1^) is the longitudinal wave speed, *f* (Hz) is the dominant frequency and *λ* (m) is the wavelength.

All signal processing steps, including FFT computation, peak detection, and wavelength estimation, were performed automatically in MATLAB using built-in functions and custom algorithms developed for this study. This approach allowed for consistent and unbiased quantification of wave propagation characteristics across different scaffolds and strain rates.

### 2.7. Statistical analysis

Statistical comparisons were performed to evaluate differences in both apparent and net mechanical properties across scaffold categories at each strain rate (n = 5 per group). A one-way analysis of variance (ANOVA) was conducted, followed by Tukey’s post hoc test to identify pairwise differences between groups (**Table S5 - S8**). The same statistical approach was applied to assess intra-group differences between rate-dependent properties within each scaffold type (**Table S9, S10**). All statistical analyses were performed using GraphPad Prism (GraphPad Software, San Diego, CA), and a significance level of *p* < 0.05 was considered statistically significant.

## 3. Results

### 3.1. Morphological Analysis

The fabricated scaffolds exhibited distinct morphological characteristics, with significant differences in fiber diameters, orientations and porosity between the PLLA/Coll and PEOT-PBT materials, as well as across the different regions of the PLLA/Coll scaffolds (**Errore. L’origine riferimento non è stata trovata.**). Scanning electron microscopy (SEM) images revealed smooth nanofibers with orientation patterns progressively transitioning from highly aligned fibers, mimicking T/L anisotropy, to more isotropic arrangements. PLLA/Coll scaffolds displayed region-specific variations in scaffold diameter (**Fig.4A-D**): the fibrocartilage-inspired (FS) region had an average of 0.72 ± 0.03 mm, the enthesis-inspired (ES) region 0.60 ± 0.07 mm, the T/L-inspired (TLS) region had an average diameter of 0.48 ± 0.04 mm. In contrast, PEOT-PBT scaffolds, designed with a uniform T/L fascicle-inspired structure (PTLS), exhibited a homogeneous cross-section of 0.61 ± 0.05 mm. Additionally, scaffolds with region-specific nanofiber diameters were fabricated, including a FS/ES region composed of randomly oriented nanofibers (nanofiber diameter: d_FS_ = 570 ± 190 nm), and a TLS consisting of randomly oriented and axially aligned nanofibers (d_TLS_ = 410 ± 130 nm). The scaffolds entirely made of PEOT-PBT exhibited nanofibers with an average diameter of 560 ± 230 nm (Errore. L’origine riferimento non è stata trovata.**Fig.4H**). Porosity measurements also confirmed structural differences among the bundle regions: PLLA/Coll scaffolds exhibited a porosity gradient, with the fibrocartilage-inspired (FS) region at 58.02 ± 8.65 %, the enthesis-inspired (ES) region at 60.99 ± 5.34 % and with the T/L-inspired (TLS) region at 50.73 ± 11.56 %. Conversely, PEOT-PBT scaffolds had a uniform porosity of 42.95 ± 9.67 %, reflecting their T/L-like structure.

Microscopic morphometric measures on microCT images (**Fig.2**, **Video S1**, **Video S2** and **Table S11**) agreed with those on SEM images, principally about trends than about absolute values. This could be expected, first because of intrinsic differences in imaging (fully 3D for microCT; 2D-3D, superficial, for SEM), second for a not controlled sample traction in microCT for avoiding micro-movements during scanning. On PLLA/Coll samples, diameter from microCT (**Fig.3A**) respect to SEM showed the same FS-ES-TLS decreasing trend, with higher absolute values; on PEOT-PBT samples (PTLS), microCT diameter was the lowest, because of maximum traction (**Fig.3B**). Moreover, sub-volume analysis (i.e., by VOI) – permitted by microCT imaging – highlighted a diameter gradient only for the PLLA/Coll samples on the ES-TLS region, confirming the gradual aspect of the realized biomimetic junction. Regarding microCT porosity (**Fig.3C-D**), it showed lower values compared to SEM – because of lower spatial resolution losing nano-porosity and of partial volume artefact masking pores in voxels that contain both material and void – but the same regional trend. MicroCT sub-volume analysis showed a gradient only for ES-TLS, transitioning from FS to TLS porosity.

**Fig. 3.**
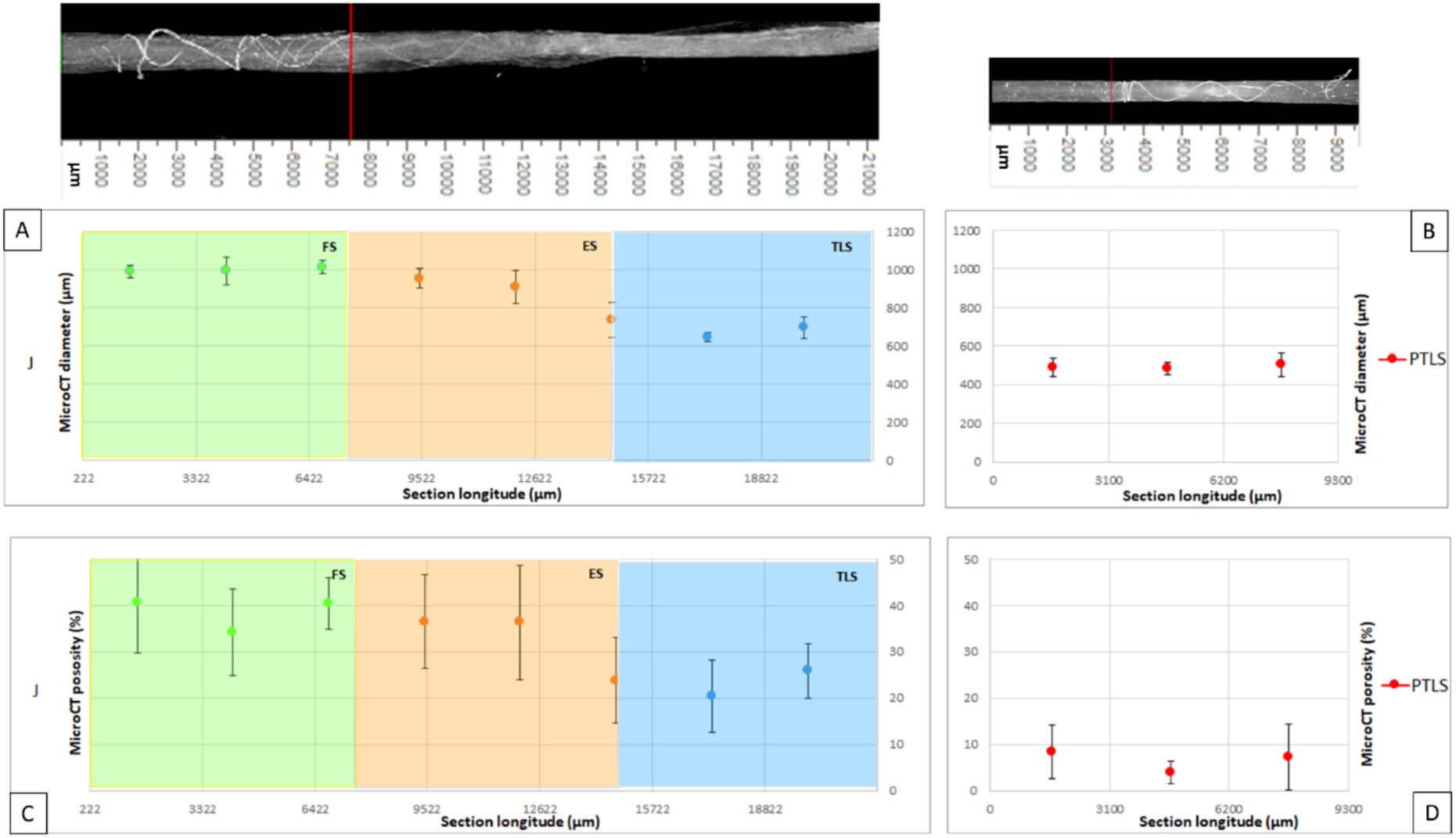
MicroCT shadow projections (above) and measure graphs (below) of PLLA/Coll (J) (left) and PEOT-PBT (PTLS) (right) samples. Diameter (A-B) and porosity (C-D) measures on microCT images were collected on longitudinal sections (2D analysis, CT-An software, Skyscan), aggregated for each VOI (shown as mean ± SD) and divided for regions (FS, ES, TLS; PTLS).

Quantitative orientation analysis (**Fig.4I**) showed an isotropic distribution of nanofibers in FS and ES samples, with only 9.8 ± 0.9% of fibers within the range 0°–6° of the main axis, and elevated proportions at higher angles, including 6.7 ± 1.2% in the range of 84°–90°. These findings indicate a reduced alignment in the FS/ES region (**Fig.4F**), which mimics the isotropic microarchitecture of the fibrocartilaginous interface. PLLA/Coll TLS samples (**Fig.4E**) exhibited a high degree of alignment, with 30.7 ± 1.9% of fibers oriented within the range 0°–6° of the scaffold axis and a markedly lower percentage 4.6 ± 2.6% at near-transverse angles 84°–90°. Similarly, PEOT-PBT bundles (**Fig.4G**) demonstrated strong axial alignment, with 30.17 ± 1.6% of fibers within the range 0°–6° and only 2.9 ± 0.9% in the 84°–90°range.

**Fig. 4.**
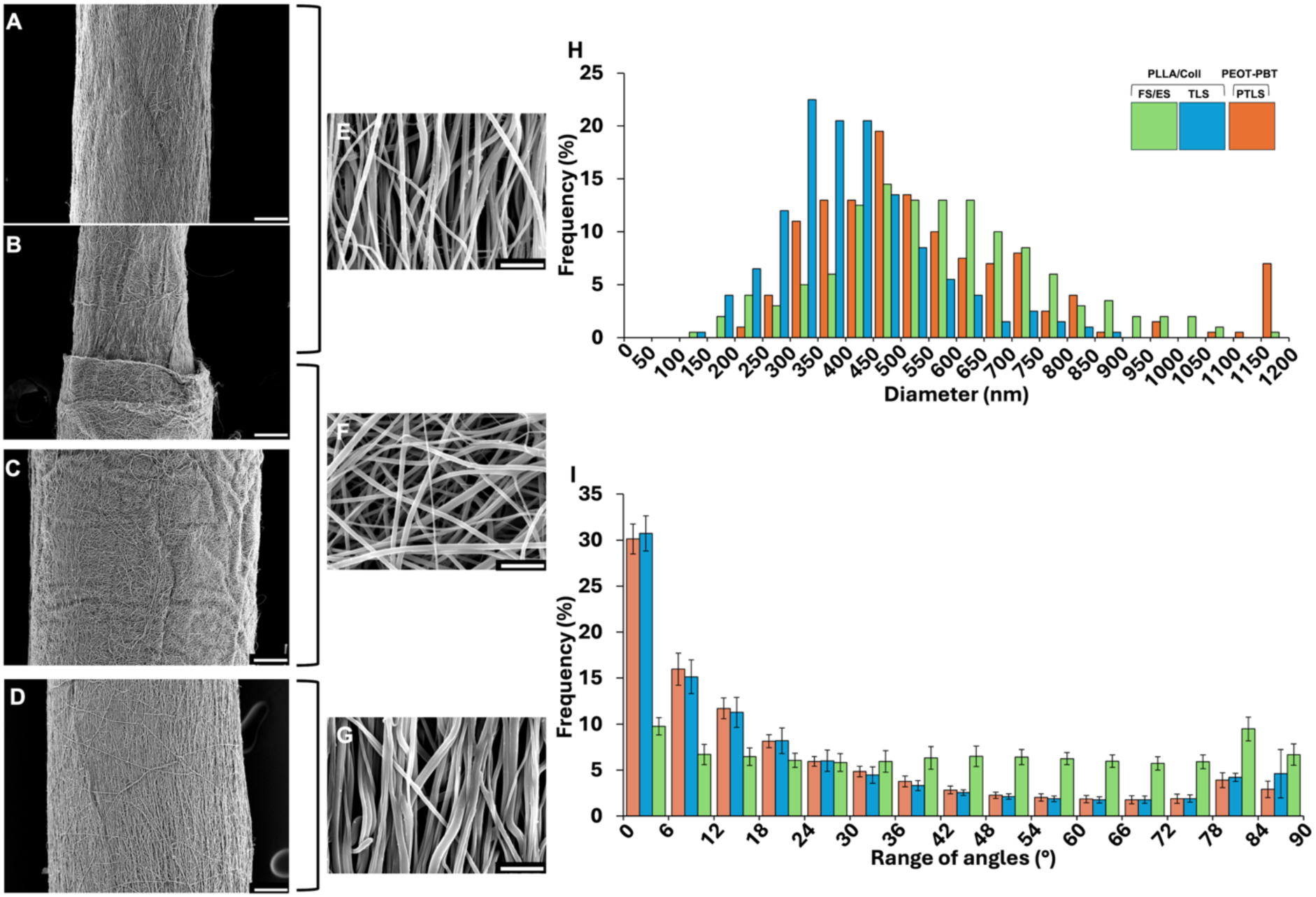
SEM morphological investigation of enthesis and T/L-inspired bundles. SEM of PLLA/Coll nanofibers (scale bar =100 µm, magnification = x150): A) T/L-inspired (TLS) side; B) enthesis-inspired (ES) side, C) fibrocartilage-inspired (FS) side. SEM of PEOT-PBT (scale bar =100 µm, magnification = x150): D) PEOT-PBT (PTLS) side; E) SEM of T/L-inspired (TLS) nanofibers (scale bar = 5 µm, magnification = x5000); F) SEM of fibrocartilage-inspired (FS)/enthesis-inspired (ES) nanofibers (scale bar = 5 µm, magnification = x5000) T/L side; G) SEM of PEOT-PBT (PTLS) nanofibers (scale bar = 5 µm, magnification = x5000) T/L side. H) Diameter distribution of the nanofibers in the different regions of the scaffolds; I) nanofibers orientation, via SEM images, on the surface of the scaffolds (0° = axial orientation; 90° = transversal orientation; images at x5000).

### 3.2 Mechanical analysis

Fibrocartilage (FS), enthesis (ES), tendon/ligament (TLS), and PEOT-PBT T/L (PTLS) inspired scaffolds were subjected to uniaxial tensile testing at multiple strain rates to characterize their mechanical behavior (Fig.5A–5C). All samples showed a ductile behaviour with large deformations with differences inter and intra PLLA/Coll and PEOT-PBT scaffolds, particularly in terms of strength, stiffness, and deformation capacity across varying geometries and strain rates (**Errore. L’origine riferimento non è stata trovata.**2). PEOT-PBT scaffolds consistently exhibited lower mechanical strength, with reduced stress and elastic modulus, when compared to their PLLA/Coll counterparts. However, they could sustain higher strains before failure (Fig.6C–6D).

**Fig. 5.**
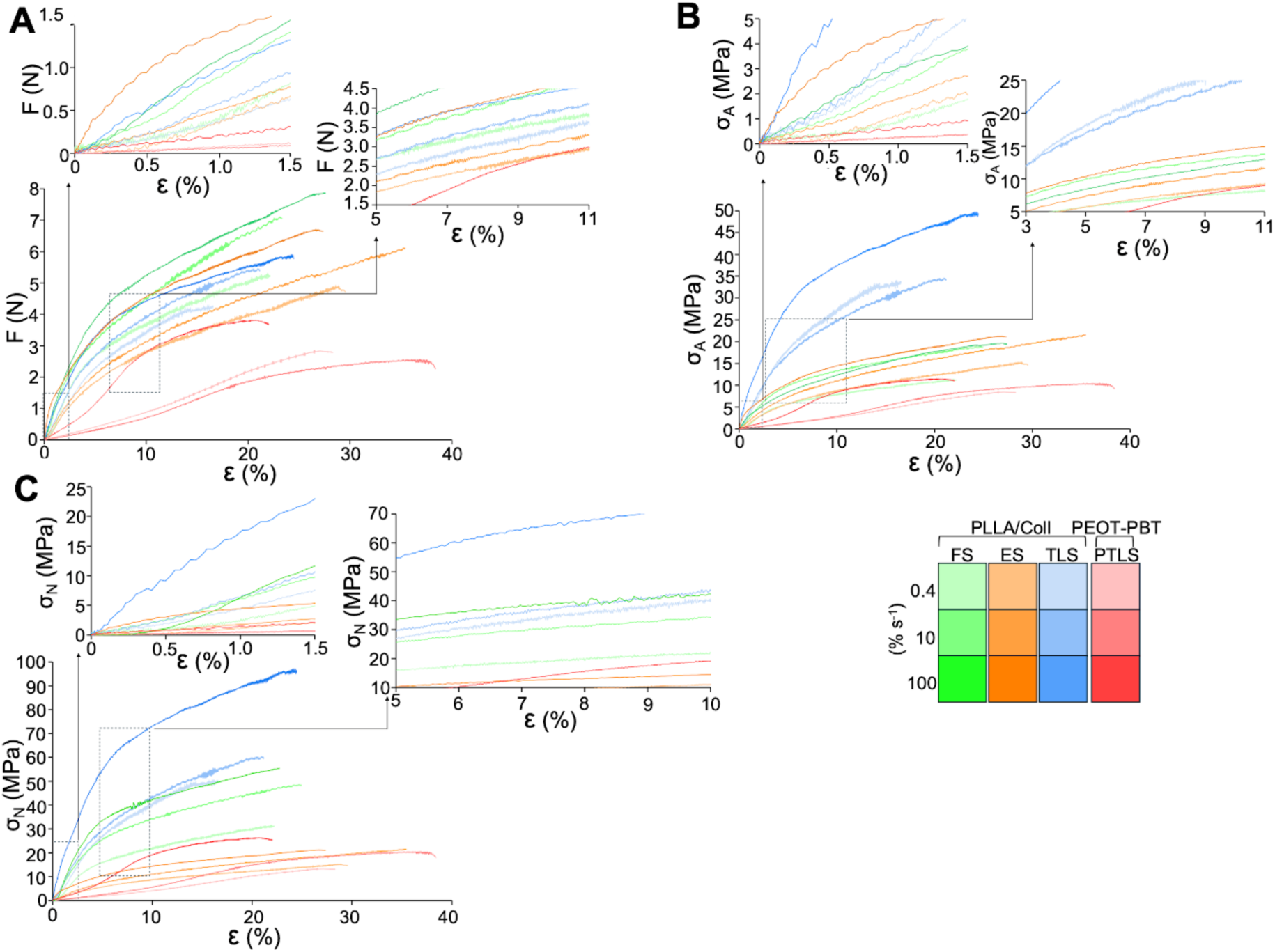
Mechanical characterization of bundles at different strain-rates (0.4, 10 and 100 % s-1): A) typical force-strain, B) apparent stress-strain and C) net stress-strain curves. T/L-inspired (TLS) scaffold enthesis-inspired (ES) scaffold, fibrocartilage-inspired (FS) scaffold, PEOT-PBT (PTLS) scaffold.

**Fig. 6.**
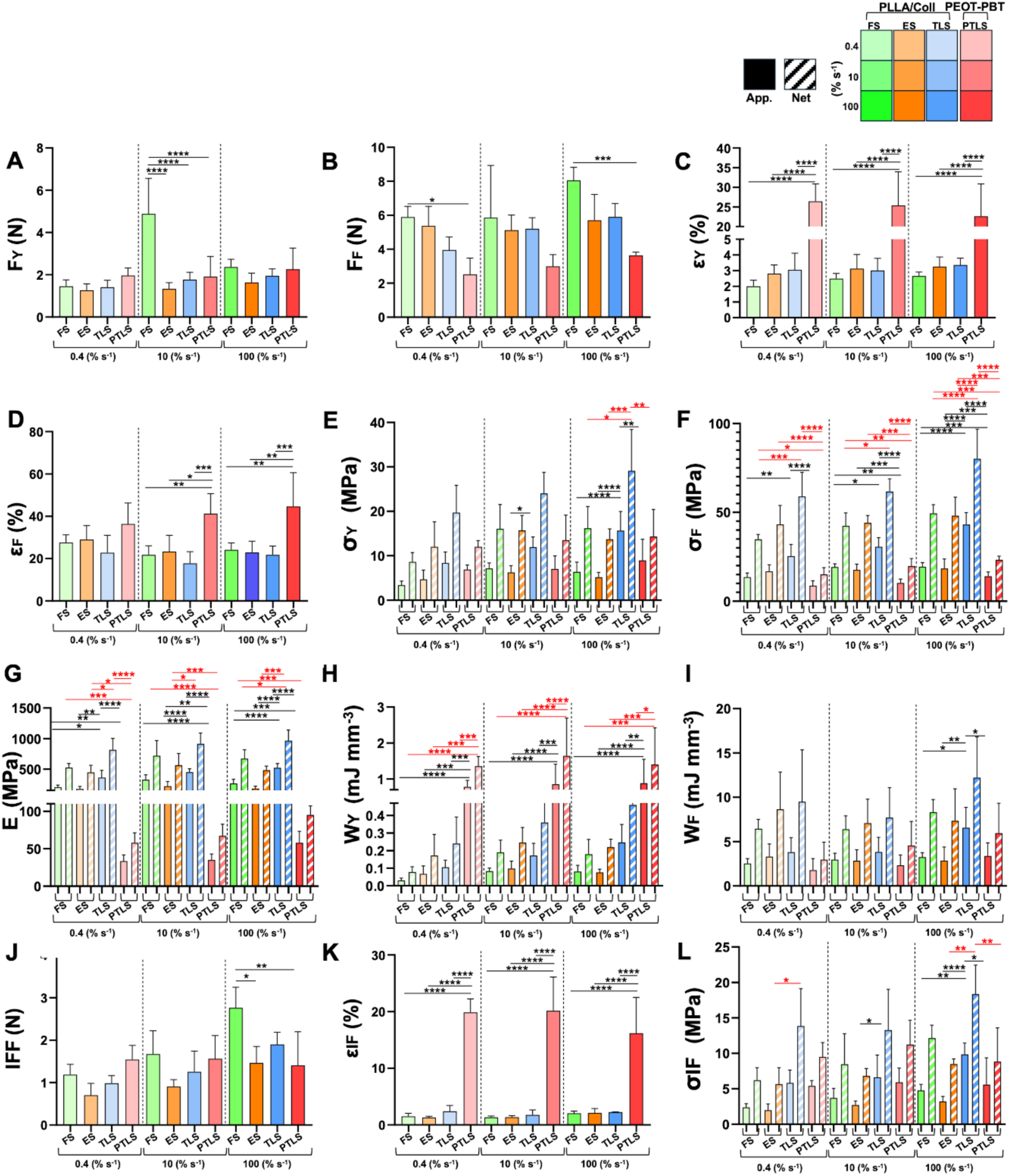
Mechanical characterization of electrospun scaffolds at different strain-rates (0.4, 10 and 100% s^-1^ apparent properties = solid lines/bars, net properties = dashed lines/bars): A) yield force (F_Y_); B) failure force (F_F_); C) yield strain (ε_Y_); D) failure strain (ε_F_); E) yield stress (σ_Y_); F) failure stress (σ_F_); G) elastic modulus (E); H) work to yield (W_Y_); I) work to failure (W_F_). Inflection point characterization of electrospun scaffolds at different strain-rates (0.4, 10 and 100 % s-1): J) inflection point force (IFF); K) inflection point strain (εIF); L) apparent and net inflection point stress (σIF). The statistically significant differences between the different bundles categories, at different strain-rates, were assessed with a one-way ANOVA followed by a Tukey post hoc (ns p>0.05, *p≤0.05, **p≤0.01, ***p≤0.001, ****p≤0.0001).The statistically significant differences between the different bundles categories, at different strain-rates, were assessed with a one-way ANOVA followed by a Tukey post hoc (ns p>0.05, *p≤0.05, **p≤0.01, ***p≤0.001, ****p≤0.0001). T/L-inspired (TLS) scaffold enthesis-inspired (ES) scaffold, fibrocartilage-inspired (FS) scaffold, PEOT-PBT (PTLS) scaffold.

Among the PLLA/Coll-based scaffolds, FS configurations reached the highest force values at yield and failure points **(Fig.6A, 6B)**. In contrast, TLS scaffolds exhibited lower force values, but showed the highest stress and stiffness, both at yield and at the failure point. At the highest strain rate (100% s⁻¹), TLS scaffolds maintained their mechanical superiority, with significantly higher yield stress and elastic modulus compared to all other configurations (Errore. L’origine riferimento non è stata trovata.**Fig.6E-6G)**. Conversely, PEOT-PBT scaffolds (PTLS) displayed lower mechanical properties across all strain rates, and they stored significant amounts of work before yield, without exhibiting a pronounced increase in energy absorption up to failure **(Fig6H, 6I)**.

No significant differences were observed in the inflection point force between PLLA/Coll and PEOT-PBT across most conditions (Fig.6J), with the exception of the FS group, which showed a higher value at 100% s^-1^ strain rate. Although PTLS bundles revealed higher inflection point strains, the inflection point stress, both apparent and net, was overall comparable between materials. Only the TLS configuration of PLLA/Coll consistently showed higher values (Fig.6K, 6L). These findings underscore fundamental differences in how each material accommodates mechanical energy and structural change under dynamic loading, informing scaffold design strategies based on targeted tissue mechanics.

### 3.3 Vibrational analysis

We investigated how the electrospun scaffolds transmitted the mechanical vibrations produced by the load applied during the uniaxial tensile test. We extracted the dominant frequency, signal amplitude at the dominant frequency, wavelength, and longitudinal wave velocity and evaluated how they are affected by the different strain rates.

The dispersion curve analysis revealed differences in the spectral behaviour of the electrospun scaffolds (**Fig.7A1-A2, Table S4**), showing an increase in dominant frequency of the transmitted mechanical waves with rising strain rate (**Fig.7B**). At low strain rates, both PLLA/Coll and PEOT-PBT exhibited similar dominant frequencies in the range 0.006–0.010 Hz. At increasing strain-rates, PEOT-PBT exhibited lower dominant frequency values compared to PLLA/Coll (0.213± 0.199 Hz and 0.539± 0.063 Hz respectively at 10 and 100% s^-1^). Conversely, PLLA/Coll showed a more pronounced increase in dominant frequency as strain rate rises.

**Fig. 7.**
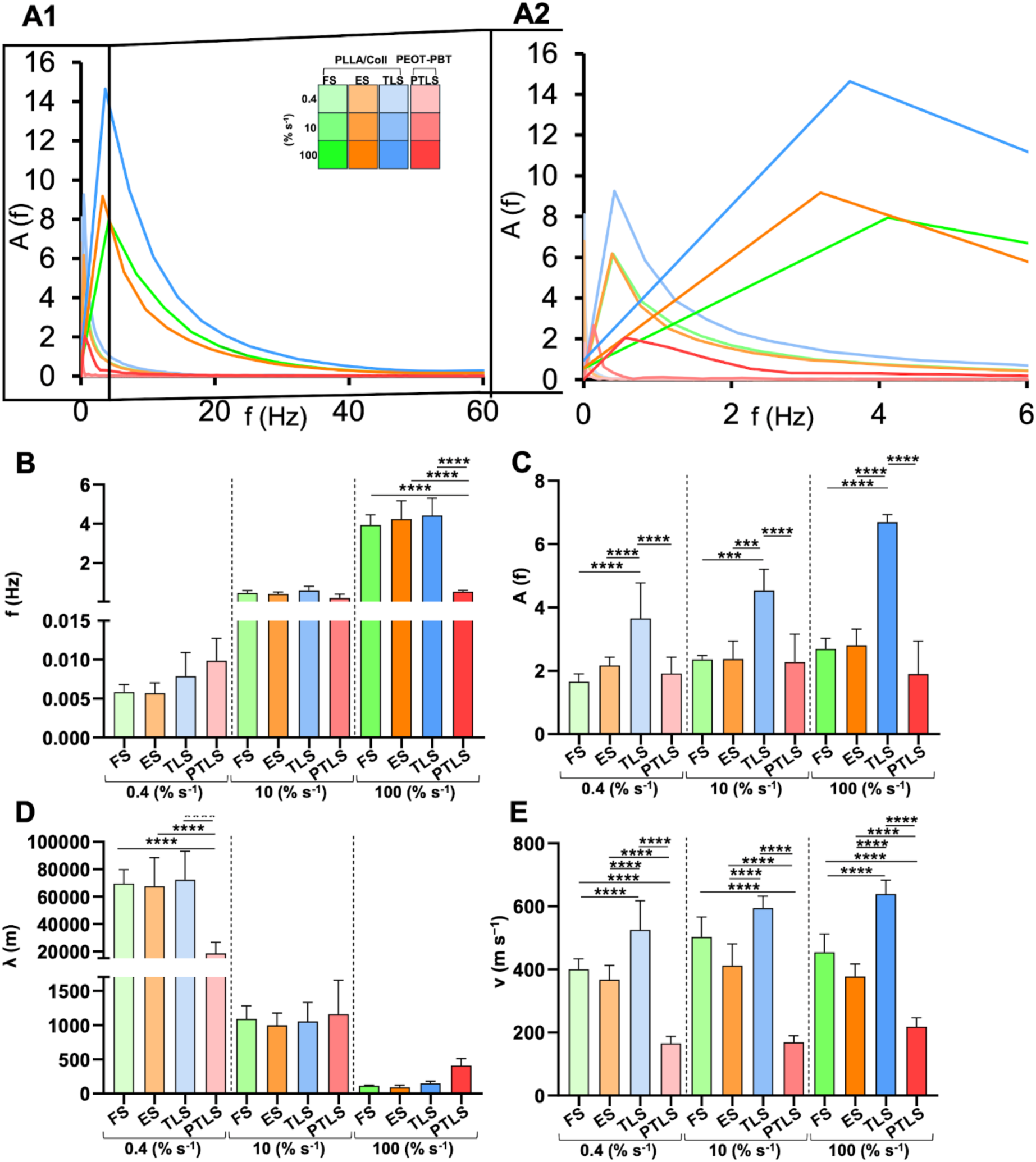
Dynamic response analysis of electrospun scaffolds at different strain-rates (0.4, 10 and 100 % s-1): A1) typical dispersion curve A2) with a zoom in the first 50 Hz, B) dominant frequency (f), C) amplitude at the dominant frequency (A); D) wavelength (λ); E) longitudinal wave speed (ν). The statistically significant differences between the different bundles categories, at different strain-rates, were assessed with a one-way ANOVA followed by a Tukey post hoc (ns p>0.05, *p≤0.05, **p≤0.01, ***p≤0.001, ****p≤0.0001). T/L-inspired (TLS) scaffold enthesis-inspired (ES) scaffold, fibrocartilage-inspired (FS) scaffold, PEOT-PBT (PTLS) scaffold.

A similar pattern was observed when analysing the amplitude at the dominant frequency. All the scaffold categories showed an increase in the signal value at the dominant frequency as the strain rate increased (Errore. L’origine riferimento non è stata trovata.**Fig.7C)**. PLLA/Coll scaffolds consistently exhibited higher peak amplitudes compared to PEOT-PBT scaffolds at all the investigated strain-rates. In contrast, PEOT-PBT scaffolds displayed a more stable amplitude profile across increasing strain rates (1.9 ± 0.5 MPa up to 2.3 ± 0.9 MPa). Overall, PLLA/Coll bundles exhibited shorter wavelengths compared to PEOT-PBT at 10 and 100% s^-1^ **(Fig.7C)**, but at increasing strain-rates both showed a decrease in wavelength. In terms of longitudinal wave velocities, at the lowest strain rate, PEOT-PBT consistently exhibited lower values (with the highest values at 166 ± 22 m s^-1^) compared to PLLA/Coll across all strain rates (**Fig.7D**).

## 4. Discussion

### 4.1 Morphological Analysis

The morphological architecture of the engineered scaffolds was designed to recapitulate the multiscale organization of the native enthesis, which comprises a spatially graded interface transitioning from aligned T/L fibers to random, mineralized fibrocartilage [7,45–48]. The PLLA/Coll scaffolds demonstrated a region-specific modulation of fiber alignment, diameter, and porosity that emulates the native hierarchical anisotropy [7,47–49]. The FS region consisted of isotropic fiber organization, the ES region exhibited progressively more isotropic fiber organization and increased porosity, consistent with the gradual structural and compositional transitions observed in the native enthesis [7,45,47,50]. Conversely, the TLS region consisted of densely packed, highly aligned fibers. This highly anisotropic architecture likely resulted from the elevated drum speed used during electrospinning, which induces preferential alignment along the main axis.

Notably, the fibrocartilage-inspired (FS) region integrated both random and aligned fibers, replicating the architectural complexity of the non-mineralized fibrocartilage including porosity gradients and intermediate fiber diameters. This offered a transition between T/L and mineralized fibrocartilage-mimetic environments, as confirmed by the microCT investigation [47,48,51]. In the enthesis-inspired (ES) region, designed to mimic the enthesis transition zone, the bundles displayed intermediate diameters (∼600 µm) and lower fiber alignment indices, resembling the histological features of non-mineralized fibrocartilage, where collagen disorganization and mineral infiltration disrupt fascicular continuity [7,45,47]. The PLLA/Coll T/L-inspired (TLS) region fibers diameter falls within the range of 90–110 μm, approximating the diameter of native T/L fascicles, which has been reported to range between 50–500 μm [36,38,49,50,52]. At the sub-bundle level, the individual fiber diameters in the electrospun scaffolds lied in the range of 0.1–1.2 μm, placing them between native collagen fibrils and collagen fibers, which can exceed 1 μm in diameter in mature T/L tissues [49,50,53]. Conversely, PEOT-PBT scaffolds (PTLS) presented a uniform morphology across the entire construct, with consistently aligned fibers and homogeneous bundle diameters of 610 μm, in line with T/L values [50].

These results demonstrate that the electrospinning process effectively modulated fiber anisotropy and diameter, allowing the scaffolds to recapitulate the architecture of the native T/L and enthesis. The spatial distribution of fiber alignment and porosity in scaffolds can influence mechanotransduction and cell fate decisions as in structurally anisotropic tissues [54–56]. MicroCT sub-volume analysis revealed sub-regional microscopic morphometric heterogeneities confined to the enthesis-inspired (ES) region, suggesting a spatially localized modulation of the scaffold mechanical response with potential implications for its dynamic behavior and the local mechanical microenvironment. Nanoscale alignment promotes tenogenic gene expression by modulating nuclear architecture and cytoskeletal tension, while disordered, porous microenvironments facilitate chondrogenic or osteogenic differentiation through diffusional and mechanical cues [53,54,55,57]. Although cellular outcomes were not explored in this work, these morphological insights offer a foundation for future studies aiming to couple localized mechanical stimulation with PEOT-PBT and PLLA/Coll scaffold-guided lineage specification [50,54,56].

### 4.2 Mechanical Analysis

The mechanical testing of the electrospun scaffolds revealed distinct mechanical profiles across the different bundle types, each designed to replicate specific zones of the T/L-to-bone interface. All samples exhibited non-linear stress-strain behaviour, including an initial toe region followed by a linear-elastic phase, consistent with the mechanical signature of native T/L and enthesis tissues [57,58].

The PLLA/Coll fibrocartilage-inspired (FS) bundles had a higher amount of material compared to other regions because of the combination of the internal layer of aligned fibers and the additional external layer of random fibers. This characteristic allowed the bundles to accommodate a higher tensile load compared to the other categories, indicating substantial load-bearing capacity.

Enthesis-inspired (ES) bundles exhibited the lowest elastic modulus (174 MPa) and the highest extensibility (strain at break of 29%) among the PLLA/Coll bundles. The hybrid fiber orientation in ES, designed to connect FS and T/L-inspired (TLS) domains, enabled progressive stress redistribution and mimicked the mechanical heterogeneity of this transitional region. ES mechanical compliance facilitates strain decoupling between the TLS and FS regions [7,47,59]. This buffering effect aligns with observations in the native enthesis, where the mineralized fibrocartilage plays a critical role in preventing stress concentrations by dissipating mechanical gradients.

The TLS PLLA/Coll bundles exhibited mechanical properties in the range of native tendon fascicles, providing a structured and directionally reinforced architecture, which can better support tendon-specific load transmission [15,36,38,55,59]. Conversely, the PEOT-PBT (PTLS) aligned bundles reflected a more compliant ductile behaviour. They exhibit lower stiffness and higher extensibility compared to the PLLA/Coll counterparts, likely due to their amorphous microstructure and higher chain mobility, suggesting more suitability for replicating the mechanics of ligamentous tissues [14,15,59]. The analysis of the inflection point in the stress-strain response further highlights the unique differences in the nonlinear mechanical behavior of the scaffolds due to their geometric arrangement.

This distinction reflects the role of scaffold architecture: the denser, more aligned fiber arrangement in TLS enhances the mechanical efficiency per cross-sectional area.

All bundle types maintained structural integrity during uniaxial loading, with no evidence of failure at the interfaces, particularly in PLLA/Coll ES constructs, confirming the mechanical continuity between regions. This is necessary for the success of multi-zonal scaffolds, as poor adhesion or mismatched mechanics across compartments are a major cause of failure in engineered enthesis grafts [47].

### 4.3 Vibrational analysis

The vibrational phenomena observed during tensile testing of electrospun T/L and enthesis fascicle-inspired bundles was systematically analysed to elucidate the influence of material and architecture on the propagation of the mechanical waves. We hypothesized that the sinusoidal mechanical signal emerging under strain is not merely due to the rupture of nanofibers, but rather reflects intrinsic oscillatory behaviours associated with bundle architecture, material stiffness, interfacial coupling and the strain rate related transfer of load.

The configuration of fibrocartilage-inspired (FS) bundles, composed of randomly oriented PLLA/Coll fibers, and of enthesis-inspired (ES) bundles, composed of randomly oriented and aligned PLLA/Coll fibers, effectively acted as mechanical dampers. Compared to PLLA/Coll T/L-inspired (TLS) bundles, FS and ES displayed reduced dominant frequency (0.006 ± 0.001 Hz), while still permitting some degree of wave propagation. Interestingly, the conical geometry of the ES bundles appeared to play a key role in modulating their wave propagation. While the aligned nanofibers within the bundles tended to promote directional transmission of vibrational energy, the tapering shape of the bundles acted as a mechanical damper. This geometric gradient disrupted the uniform propagation of mechanical waves, dissipating energy and effectively reducing the stiffness typically associated with aligned fiber systems. As a result, the frequency-dependent mechanical behaviour of the ES bundles shifted toward a more fibrocartilage-like profile. This behavior highlights the role of interfacial regions in decoupling mechanical environments, preventing direct propagation of strain-induced oscillations across functionally distinct domains. In native entheses, this mechanical discontinuity is crucial for minimizing shear stress concentrations and supporting spatially regulated cellular differentiation [60,61].

T/L-inspired (TLS) and PTLS aligned bundles were both designed to mimic the native T/L fascicles, but they exhibited markedly distinct vibrational profiles. PLLA/Coll TLS bundles showed high-wave transmission, with velocities reaching 639 ± 44 m s^-1^, and elevated wave amplitudes. These features resemble the mechanical behavior of natural tendon fascicles and they arise from the semi-crystalline structure of PLLA, which contributes to the directional wave propagation [15,61,62]. In contrast, PEOT-PBT, despite also being aligned, is characterized by a lower degree of crystallinity and higher elasticity, thus it exhibited attenuated vibrational responses, with lower amplitudes and wave velocity, which did not exceed 218 ± 29 m s^-1^. The elastomeric nature of the polymer raises from combination of soft PEOT and hard PBT blocks, leading to reduced wave transmission and increased energy dissipation [64,65]. Interestingly, although both materials were designed to mimic the T/L structure, the TLS bundles exhibited a vibration transmission behavior more consistent with tendon function, suggesting greater potential to replicate tendon-like mechanical cues. In contrast, the PEOT-PBT samples displayed a more elastic response, indicative of mechanical characteristics typically associated with ligamentous tissues.

The differential wave transmission observed across the bundle types holds profound implications for T/L-to-bone tissue engineering [66]. Mechanical stimulation, particularly in the form of high-frequency, low-amplitude vibrations, promotes osteogenic and chondrogenic differentiation of mesenchymal stem cells (MSCs) when applied in a frequency- and amplitude-dependent manner [67–69]. Aligned, high-stiffness scaffolds such as PLLA/Coll T/L-inspired (TLS) may thus favour stem cells differentiation via directional mechanical cues, while more damped regions such as fibrocartilage-inspired (FS) and enthesis-inspired (ES) may promote a chondrogenic or osteogenic fate due to their attenuated, diffused mechanical environments [69]. Excessively low vibration transmission, as observed in PEOT-PBT, can inhibits mechanoresponsive pathways, potentially advantageous for anti-inflammatory or protective contexts, but suboptimal where active mechano-induction is desired.

The observed differences in mechanical wave propagation are consistent with the hierarchical organization of native T/L tissue, where collagen fibril diameter, fascicle alignment, and interfibrillar matrix composition all contribute to region-specific mechanical responses [60]. The scaffolds did not rely on externally imposed oscillations; mechanical loading elicited intrinsic vibrational responses. These were driven by viscoelastic, strain-rate-dependent transmission, arising from both progressive nanofiber rupture and the gradual build-up of scaffold tension, and further shaped by scaffold geometry. The PLLA/Coll TLS bundles showed the highest capacity to transmit mechanical waves, followed by the PLLA/Coll FS and ES configurations, which exhibited intermediate transmission, while the PEOT-PBT samples displayed the lowest transmission capacity. Conversely, when considering the ability to attenuate mechanical vibrations, the PEOT-PBT scaffolds were the most effective, followed by the PLLA/Coll FS and ES bundles, with the PLLA/Coll TLS bundles showing the least damping behavior. The trends of the two scaffolds reflect the combined effects of material properties and scaffold architecture, providing insights into how composition and fibre organization jointly influence vibrational behaviour across varying strain rates. These findings underscore the relevance of scaffold geometry and composition in shaping dynamic mechanical responses and suggest that different material configurations may be more suitable for tendon or ligament regeneration depending on whether vibrational transmission or damping is desired.

## 5. Conclusions

This study describes the mechanical stimulation-induced vibrational response of biomaterials for tendon/ligament (T/L) and enthesis regeneration. Electrospun PLLA/Coll and PEOT-PBT scaffolds were designed to mimic native T/L tissue and its bone interface. SEM and micro-CT confirmed that fiber dimensions and orientations closely reproduce those of the native ECM, while tensile tests showed comparable mechanical properties, supporting PEOT-PBT for ligament and PLLA/Coll for tendon and enthesis regeneration. Vibrational behavior under tensile loading was assessed using Fast Fourier Transform (FFT), identifying architecture and material-specific vibrational signatures associated with mechanical stimulation.

By combining mechanical testing and frequency-domain analysis, this study provides new insights in how scaffold composition and architecture influence the transmission of mechanical signals across engineered T/L-to-bone constructs. PEOT-PBT scaffolds generally attenuate mechanical signals across all tested strain rates, while PLLA/Coll scaffolds promote greater propagation, particularly with aligned nanofibers. Notably, the transition zone in PLLA/Coll scaffolds, connecting aligned and randomly aligned nanofibers, reduces signal propagation acting as a mechanical decoupler. These findings suggest that such differences in mechanical cues transmission could shape tissue and cell responses across graded scaffold designs.

Building on this, future work will examine how these vibrational properties influence cell differentiation under mechanical stimulation, testing the functional link between scaffold microstructure, mechanical signal transmission, and cell fate. This framework lays the groundwork for designing biomaterials with controlled mechanical microenvironments to direct T/L and enthesis regeneration.

## 6. Acknowledgments

Horizon Europe Marie Skłodowska Curie Postdoctoral Fellowship 3NTHESES (n.101061826) is greatly acknowledged for funding the work. Type I collagen was kindly provided by Kensey Nash Corporation d/b/a DSM Biomedical (Exton, USA).

## 7. CRediT authorship contribution statement

**A. Di Lorenzo:** Writing – original draft, Methodology, Investigation, Visualization, Formal analysis, Data curation. **T. Ten Brink:** Writing – original draft, Investigation, Formal analysis. **G. Marchiori:** Writing – original draft, Methodology, Investigation, Visualization, Formal analysis, Data curation. **G. Giavaresi:** Writing – review & editing, Resources, Supervision. **L. Moroni:** Writing – review & editing, Methodology, Resources, Funding acquisition, Project administration, Conceptualization, Supervision. **M. van Griensven:** Writing – review & editing, Methodology, Resources, Funding acquisition, Project administration, Conceptualization, Supervision. **A. Sensini:** Writing – original draft, Methodology, Investigation, Visualization, Resources, Funding acquisition, Formal analysis, Data curation, Project administration, Conceptualization, Supervision.

## 8. Declaration of competing interest

The authors declare that they have no known competing financial interests or personal relationships that could have appeared to influence the work reported in this paper.

## 9. Data availability

Data will be made available from the authors under request.

## 10.

## Appendix A. Supplementary Data

Supplementary material of this article can be found online.

## Notes

### Competing Interest Statement

The authors have declared no competing interest.

