## Supplementary_Material for "Highlighting strain rate dependent vibrational behavior of electrospun bundles for tendon/ligament and enthesis fascicle tissue regeneration"

**File include:**
Video: Video S1, Video S2

Table: Table S1, S2, S3, S4, S5, S6, S7, S8, S9, S10, S11

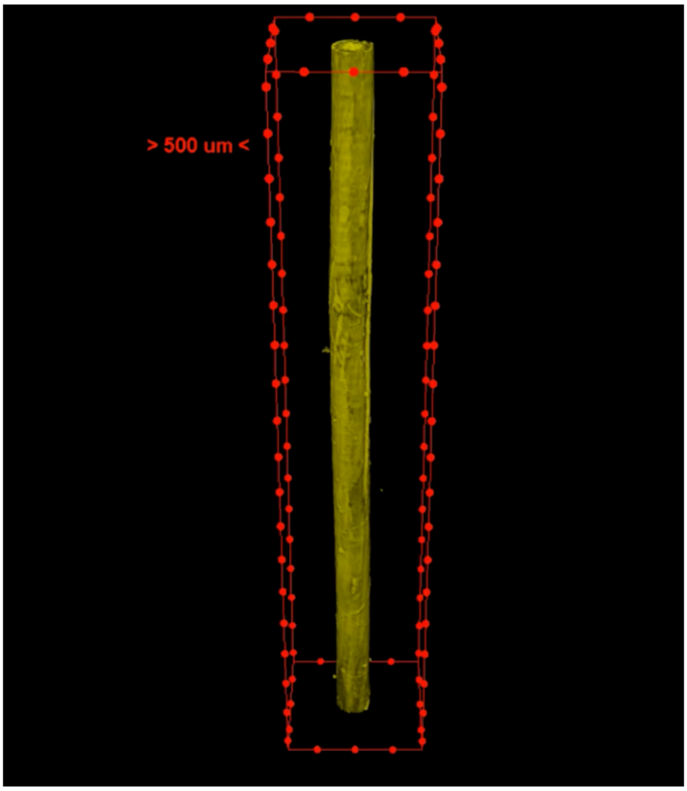


**Video S1.** Micro-CT movie of a PEOT-PBT bundle.

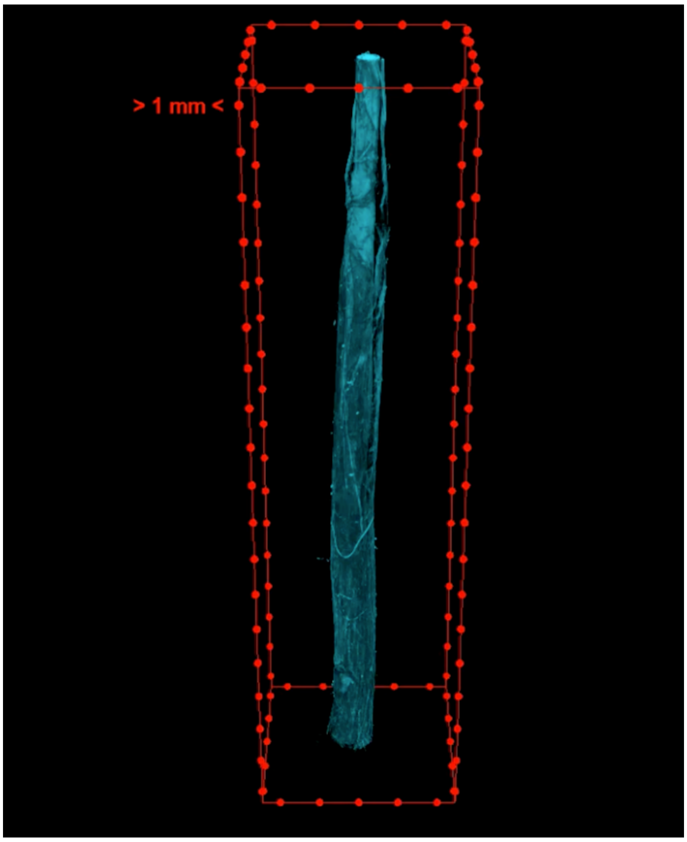


**Video S2.** Micro-CT movie of a PLLA/Coll junction.

**Table S1.** Mean morphological parameters of the different categories of scaffolds.

| Sample | l [mm] | $l_0$ [mm] | d [µm] | D [mm] | A [mm$^2$] | υ [-] | P [%] | W [mg] | $w_0$ [mg] |
|---|---|---|---|---|---|---|---|---|---|

| | | | | | | | | | |
|---|---|---|---|---|---|---|---|---|---|
| FS | 40 | 20 | 0.57 ± 0.19 | 0.72 ± 0.03 | 0.42 ± 0.03 | 0.4 ± 0.1 | 57.8 ± 8.7 | 7.3 ± 1.2 | 4.4 ± 0.7 |
| ES | 40 | 20 | | 0.60 ± 0.07 | 0.29 ± 0.02 | 0.4 ± 0.1 | 60.9 ± 5.0 | 4.9 ± 0.9 | 3.1 ± 0.6 |
| TLS | 40 | 20 | 0.41 ± 0.13 | 0.48 ± 0.04 | 0.16 ± 0.03 | 0.5 ± 0.1 | 50.7 ± 11.6 | 3.3 ± 0.6 | 1.9 ± 0.3 |
| PTLS | 40 | 20 | 0.56 ± 0.23 | 0.61 ± 0.05 | 0.29 ± 0.05 | 0.6 ± 0.1 | 43.0 ± 9.7 | 5.8 ± 1.1 | 3.7 ± 0.7 |

**Table S2.** Apparent and net mechanical properties of scaffolds at the different strain-rates investigated.

| | Strain-rate [% $s^{-1}$] | Sample | $F_y$ [N] | $F_F$ [N] | $\varepsilon_Y$ [%] | $\varepsilon_F$ [%] | $\sigma_Y$ [MPa] | $\sigma_F$ [MPa] | E [MPa] | $W_y$ [mJ $mm^{-3}$] | $W_F$ [mJ $mm^{-3}$] |
|---|---|---|---|---|---|---|---|---|---|---|---|
| **App** | 0.4 | FS | 1.5 ± 0.3 | 5.9 ± 0.6 | 2.0 ± 0.4 | 27.6 ± 3.6 | 3.4 ± 0.9 | 13.6 ± 2.3 | 204.8 ± 34.6 | 0.3 ± 0.1 | 2.5 ± 0.6 |
| | | ES | 1.3 ± 0.3 | 5.4 ± 1.1 | 2.8 ± 0.5 | 29.0 ± 6.7 | 4.7 ± 2.1 | 16.8 ± 3.6 | 173.5 ± 43.7 | 0.7 ± 0.5 | 3.3 ± 1.4 |
| | | TLS | 1.4 ± 0.3 | 4.0 ± 0.7 | 3.1 ± 1.1 | 22.7 ± 0.4 | 8.4 ± 2.5 | 25.4 ± 6.7 | 359.9 ± 116.1 | 0.1 ± 0.0 | 3.8 ± 1.7 |
| | | PTLS | 2.0 ± 0.4 | 2.5 ± 1.0 | 26.4 ± 4.4 | 36.4 ± 9.8 | 6.9 ± 1.0 | 8.8 ± 2.8 | 33.5 ± 8.4 | 0.8 ± 0.2 | 1.8 ± 1.3 |
| | 10 | FS | 4.9 ± 1.7 | 5.9 ± 3.1 | 2.5 ± 0.3 | 21.7 ± 4.3 | 3.4 ± 0.9 | 19.3 ± 1.7 | 325.3 ± 81.5 | 0.1 ± 0.0 | 3.0 ± 0.7 |
| | | ES | 1.3 ± 0.3 | 5.1 ± 0.9 | 3.1 ± 0.9 | 23.4 ± 7.7 | 6.2 ± 1.5 | 17.6 ± 3.2 | 221.0 ± 75.2 | 0.1 ± 0.0 | 2.9 ± 1.3 |
| | | TLS | 1.8 ± 0.4 | 5.2 ± 0.7 | 3.0 ± 0.8 | 17.8 ± 5.5 | 11.9 ± 2.3 | 30.7 ± 5.0 | 450.4 ± 55.9 | 0.2 ± 0.1 | 3.8 ± 1.7 |
| | | PTLS | 1.9 ± 1.0 | 3.0 ± 0.7 | 25.4 ± 8.5 | 41.3 ± 9.5 | 7.1 ± 2.9 | 10.3 ± 2.2 | 35.1 ± 7.8 | 0.9 ± 0.6 | 2.3 ± 1.2 |
| | 100 | FS | 2.4 ± 0.4 | 8.1 ± 0.7 | 2.7 ± 0.3 | 24.2 ± 3.3 | 6.4 ± 2.2 | 19.5 ± 2.3 | 265.7 ± 67.0 | 0.1 ± 0.0 | 3.3 ± 0.5 |
| | | ES | 1.6 ± 0.4 | 5.7 ± 1.5 | 3.3 ± 0.6 | 22.9 ± 5.4 | 5.2 ± 1.1 | 18.4 ± 5.4 | 182.7 ± 37.1 | 0.1 ± 0.0 | 2.9 ± 1.5 |
| | | TLS | 1.9 ± 0.3 | 5.9 ± 0.8 | 3.4 ± 0.4 | 21.7 ± 4.2 | 15.7 ± 4.3 | 43.2 ± 6.8 | 521.0 ± 69.9 | 0.2 ± 0.1 | 6.6 ± 2.3 |
| | | PTLS | 2.3 ± 1.0 | 3.6 ± 0.2 | 22.6 ± 8.2 | 44.6 ± 15.9 | 9.0 ± 4.7 | 14.0 ± 2.4 | 57.9 ± 15.3 | 0.9 ± 0.7 | 3.4 ± 1.5 |
| **Net** | 0.4 | FS | - | - | - | - | 8.7 ± 2.1 | 34.7 ± 2.8 | 523.4 ± 68.9 | 0.1 ± 0.0 | 6.5 ± 1.0 |
| | | ES | - | - | - | - | 12.1 ± 5.6 | 43.3 ± 10.6 | 444.5 ± 117.7 | 0.2 ± 0.1 | 8.6 ± 4.2 |
| | | TLS | - | - | - | - | 19.7 ± 6.1 | 59.0 ± 13.5 | 815.9 ± 186.6 | 0.3 ± 0.1 | 9.5 ± 5.8 |
| | | PTLS | - | - | - | - | 12.0 ± 1.4 | 15.1 ± 3.8 | 58.0 ± 13.3 | 1.4 ± 0.3 | 3.0 ± 2.0 |
| | 10 | FS | - | - | - | - | 16.1 ± 5.5 | 42.4 ± 7.3 | 719.7 ± 248.8 | 0.2 ± 0.1 | 6.4 ± 1.5 |
| | | ES | - | - | - | - | 15.7 ± 3.4 | 44.1 ± 4.1 | 563.9 ± 193.2 | 0.2 ± 0.1 | 7.1 ± 2.7 |
| | | TLS | - | - | - | - | 24.0 ± 4.7 | 61.7 ± 7.1 | 916.1 ± 175.3 | 0.3 ± 0.1 | 3.8 ± 1.7 |
| | | PTLS | - | - | - | - | 13.5 ± 5.6 | 19.7 ± 4.3 | 67.2 ± 15.5 | 1.6 ± 1.1 | 4.6 ± 2.7 |
| | 100 | FS | - | - | - | - | 16.2 ± 4.9 | 49.4 ± 4.9 | 672.7 ± 248.8 | 0.2 ± 0.1 | 8.3 ± 1.4 |
| | | ES | - | - | - | - | 13.7 ± 2.3 | 48.2 ± 10.4 | 481.9 ± 68.0 | 0.2 ± 0.0 | 7.4 ± 3.6 |
| | | TLS | - | - | - | - | 29.1 ± 9.3 | 80.1 ± 16.6 | 966.4 ± 175.9 | 0.5 ± 0.2 | 12.3 ± 4.8 |
| | | PTLS | - | - | - | - | 16.2 ± 4.9 | 49.4 ± 4.9 | 672.7 ± 248.8 | 0.2 ± 0.1 | 8.3 ± 1.4 |

**Table S3.** Apparent and net inflection point properties of scaffolds at the different strain-rates investigated.

| Strain-rate [% $s^{-1}$] | Sample | IFF [N] | IF$\varepsilon$ [%] | IF$\sigma_{APP}$ [MPa] | IF$\sigma_{NET}$ [MPa] |
|---|---|---|---|---|---|
| 0.4 | FS | 1.2 ± 0.2 | 1.5 ± 0.5 | 2.4 ± 0.5 | 6.2 ± 1.8 |
| | ES | 1.3 ± 0.7 | 1.6 ± 0.5 | 2.7 ± 1.3 | 6.4 ± 3.4 |
| | TLS | 1.0 ± 0.2 | 2.4 ± 1.0 | 5.8 ± 1.8 | 13.8 ± 5.3 |
| | PTLS | 1.5 ± 0.3 | 19.9 ± 2.4 | 5.4 ± 0.8 | 9.5 ± 2.3 |
| 10 | FS | 1.7 ± 0.6 | 1.3 ± 0.3 | 3.7 ± 1.3 | 8.5 ± 4.3 |
| | ES | 0.9 ± 0.2 | 1.4 ± 0.3 | 2.7 ± 0.6 | 6.8 ± 1.0 |
| | TLS | 1.3 ± 0.5 | 1.8 ± 0.9 | 6.6 ± 3.1 | 13.3 ± 5.8 |
| | PTLS | 1.6 ± 0.5 | 20.2 ± 5.9 | 5.9 ± 2.0 | 11.3 ± 3.4 |
| 100 | FS | 2.8 ± 0.5 | 2.1 ± 0.4 | 4.8 ± 0.8 | 12.1 ± 1.8 |
| | ES | 1.5 ± 0.4 | 2.1 ± 0.8 | 3.2 ± 0.7 | 8.5 ± 0.8 |
| | TLS | 1.9 ± 0.3 | 2.3 ± 0.1 | 9.9 ± 1.6 | 18.4 ± 4.1 |
| | PTLS | 1.4 ± 0.8 | 16.2 ± 6.2 | 5.6 ± 3.8 | 8.8 ± 4.7 |

**Table S4.** Frequency-dependent properties of scaffolds across tested strain-rates.

| Strain-rate [% $s^{-1}$] | Sample | f [Hz] | A [MPa] | $\lambda$ [m] | v [m $s^{-1}$] |
|---|---|---|---|---|---|
| 0.4 | FS | 0.006 ± 0.001 | 1.7 ± 0.2 | 69548 ± 10096 | 400 ± 33 |
| | ES | 0.006 ± 0.001 | 2.2 ± 0.3 | 67583 ± 20804 | 367 ± 45 |
| | TLS | 0.007 ± 0.003 | 3.6 ± 1.1 | 72370 ± 1.8 | 526 ± 92 |
| | PTLS | 0.010 ± 0.003 | 1.9 ± 0.5 | 18609 ± 8091 | 166 ± 22 |
| 10 | FS | 0.5 ± 0.1 | 2.4 ± 0.1 | 1091 ± 191 | 503 ± 64 |
| | ES | 0.4 ± 0.1 | 2.4 ± 0.6 | 996 ± 183 | 413 ± 68 |
| | TLS | 0.6 ± 0.2 | 4.5 ± 1.3 | 1055 ± 275 | 13.3 ± 5.8 |
| | PTLS | 0.213± 0.199 | 2.3 ± 0.9 | 1159 ± 496 | 170 ± 19 |
| 100 | FS | 3.9 ± 0.5 | 2.7 ± 0.3 | 115 ± 9.2 | 454 ± 58 |
| | ES | 4.2 ± 0.9 | 2.8 ± 0.5 | 94 ± 29 | 377 ± 39 |
| | TLS | 1.9 ± 0.3 | 6.7 ± 0.2 | 150 ± 33 | 639 ± 44 |
| | PTLS | 0.539 ± 0.063 | 1.9 ± 1.0 | 413 ± 102 | 218 ± 29 |

**Table S5**. The significance of differences between the different categories of scaffolds in terms of apparent and net tensile mechanical properties assessed with a one-way ANOVA followed by a Tukey post hoc (ns p>0.05, *p≤0.05, **p≤0.01, ***p≤0.001, ****p≤0.0001).

| | Strain-rate [% $s^{-1}$] | Sample | $F_y$ [N] | $F_F$ [N] | $\varepsilon_Y$ [%] | $\varepsilon_F$ [%] | $\sigma_Y$ [MPa] | $\sigma_F$ [MPa] | E [MPa] | $W_y$ [mJ $mm^{-3}$] | $W_F$ [mJ $mm^{-3}$] |
|---|---|---|---|---|---|---|---|---|---|---|---|
| **App** | 0.4 | FS vs. ES | ns | ns | ns | ns | ns | ns | ns | ns | ns |
| | | FS vs. TLS | ns | ns | ns | ns | ns | ** (p=0.0019) | ** (p=0.0080) | ns | ns |
| | | FS vs. PTLS | **** (p<0,0001) | ** (p=0.0035) | **** (p<0,0001) | ns | ns | ns | ** (p=0.0022) | ** (p=0.0017) | ns |
| | | ES vs. TLS | ns | ns | ns | ns | ns | ns | *** (p=0.0006) | ns | ns |
| | | ES vs. PTLS | **** (p<0,0001) | ** (p=0.0249) | **** (p<0,0001) | ns | ns | ns | * (p=0.0251) | ** (p=0.0036) | ns |
| | | TLS vs. PTLS | **** (p<0,0001) | | **** (p<0,0001) | | | **** (p<0,0001) | **** (p<0,0001) | ** (p=0.0073) | ns |
| **Net** | 0.4 | FS vs. ES | - | - | - | - | ns | ns | ns | ns | ns |
| | | FS vs. TLS | - | - | - | - | ns | ** (p=0,0024) | ns | ns | ns |
| | | FS vs. PTLS | - | - | - | - | ns | * (p=0,0279) | *** (p=0.0002) | ** (p=0.0018) | ns |
| | | ES vs. TLS | - | - | - | - | ns | ns | ** (p=0.0069) | ns | ns |
| | | ES vs. PTLS | - | - | - | - | ns | *** (p=0,0002) | ** (p=0.0041) | ** (p=0.0051) | ns |
| | | TLS vs. PTLS | - | - | - | - | ns | **** (p<0,0001) | **** (p<0,0001 | * (p=0.0105) | ns |
| **App** | 10 | FS vs. ES | ns | ns | ns | ns | ns | ns | ns | ns | ns |
| | | FS vs. TLS | ns | ns | ns | ns | ns | ** (p=0,0029) | ns | ns | ns |
| | | FS vs. PTLS | ns | ns | **** (p<0,0001) | * (p=0,0117) | ns | * (p=0,0411) | **** (p<0,0001) | ** (p=0.0012) | ns |
| | | ES vs. TLS | ns | ns | **** (p<0,0001) | ns | ns | *** (p=0,0003) | **** (p<0,0001) | ns | ns |
| | | ES vs. PTLS | ns | ns | **** (p<0,0001) | * (p=0,0294) | ns | ns | ns *** (p=0,0006) | ** (p=0.0016) | ns |
| | | TLS vs. PTLS | ns | ns | **** (p<0,0001) | *** (p=0,0001) | ns | **** (p<0,0001) | **** (p<0,0001) | ** (p=0.0065) | ns |
| **Net** | 10 | FS vs. ES | - | - | - | - | ns | ns | ns | ns | ns |
| | | FS vs. TLS | - | - | - | - | ns | * (p=0,0340) | ns | ns | ns |
| | | FS vs. PTLS | - | - | - | - | ns | ** (p=0,0056) | **** (p<0,0001) | ns | ns |
| | | ES vs. TLS | - | - | - | - | ns | ns | * (p<0,0129) | ns | ns |
| | | ES vs. PTLS | - | - | - | - | ns | ** (p=0,0021) | **** (p<0,0001) | *** (p=0.0005) | ns |
| | | TLS vs. PTLS | - | - | - | - | ns | **** (p<0,0001) | **** (p<0,0001) | ns | ns |
| **App** | 100 | FS vs. ES | ns | ns | ns | ns | ns | ns | ns | ns | ns |
| | | FS vs. TLS | ns | ns | ns | ns | ns | **** (p<0,0001) | **** (p<0,0001) | ns | ns |
| | | FS vs. PTLS | ns | ns | **** (p<0,0001) | ** (p=0,0067) | ns | ns | **** (p<0,0001) | *** (p=0.0006) | ns |
| | | ES vs. TLS | ns | ns | ns | ns | ns | **** (p<0,0001) | **** (p<0,0001) | ns | ns |
| | | ES vs. PTLS | ns | ns | **** (p<0,0001) | ** (p=0,0030) | ns | ns | ns | *** (p=0.0005) | ns |
| | | TLS vs. PTLS | ns | ns | **** (p<0,0001) | ** (p=0,00015) | ns | **** (p<0,0001) | **** (p<0,0001) | * (p=0.0139) | ns |

| | | | $F_y$ | $F_F$ | $\varepsilon_Y$ | $\varepsilon_F$ | $\sigma_Y$ | $\sigma_F$ | E | $W_y$ | $W_F$ |
|---|---|---|---|---|---|---|---|---|---|---|---|
| **Net** | 100 | FS vs. ES | - | - | - | - | ns | ns | ns | ns | ns |
| | | FS vs. TLS | - | - | - | - | ns | **** (p<0,0001) | ns | ns | ns |
| | | FS vs. PTLS | - | - | - | - | ns | *** (p=0,0008) | **** (p<0,0001) | ** (p=0.0032) | ns |
| | | ES vs. TLS | - | - | - | - | ** (p=0,0012) | **** (p<0,0001) | *** (p=0,0001) | ns | ns |
| | | ES vs. PTLS | - | - | - | - | ns | ** (p=0,0016) | ** (p<0,0041) | ** (p=0.0049) | ns |
| | | TLS vs. PTLS | - | - | - | - | ** (p=0,0023) | **** (p<0,0001) | **** (p<0,0001) | ns | ns |

**Table S6.** The significance of differences of apparent and net mechanical properties of bundles at different strain-rates assessed with a one-way ANOVA followed by a Tukey post hoc (ns p>0.05, *p≤0.05, **p≤0.01, ***p≤0.001, ****p≤0.0001).

| | Sample | $F_y$ [N] | $F_F$ [N] | $\varepsilon_Y$ [%] | $\varepsilon_F$ [%] | $\sigma_Y$ [MPa] | $\sigma_F$ [MPa] | E [MPa] | $W_y$ [mJ mm$^{-3}$] | $W_F$ [mJ mm$^{-3}$] |
|---|---|---|---|---|---|---|---|---|---|---|
| **App** | FS 0.4 vs 10 | ns | ns | ns | ns | ns | ns | ns | ns | ns |
| | FS 0.4 vs 100 | **** (p<0,0001) | ns | ns | ns | ns | ns | ns | ns | ns |
| | FS 10 vs 100 | ns | ns | ns | ns | ns | ns | ns | ns | ns |
| | ES 0.4 vs 10 | **** (p<0,0001) | ns | ns | ns | ns | ns | ns | ns | ns |
| | ES 0.4 vs 100 | ns | ns | ns | ns | ns | ns | ns | ns | ns |
| | ES 10 vs 100 | ns | ns | ns | ns | ns | ns | ns | ns | ns |
| | TLS 0.4 vs 10 | ns | ns | ns | ns | ns | ns | ns | ns | ns |
| | TLS 0.4 vs 100 | ns | ns | ns | ns | ** (p=0,0022) | ns | ** (p=0,0050) | ns | ns |
| | TLS 10 vs 100 | ns | ns | ns | ns | ns | *** (p=0,0008) | ns | ns | ns |
| | PTLS 0.4 vs 10 | ns | ns | ns | ns | ns | ns | ns | ns | ns |
| | PTLS 0.4 vs 100 | ns | ns | ns | ns | ns | ns | ns | ns | ns |
| | PTLS 10 vs 100 | ns | ns | ns | ns | ns | ns | ns | ns | ns |
| **Net** | FS 0.4 vs 10 | - | - | - | - | ns | ns | ns | ns | ns |
| | FS 0.4 vs 100 | - | - | - | - | ns | ns | ns | ns | ns |
| | FS 10 vs 100 | - | - | - | - | ns | ns | ns | ns | ns |
| | ES 0.4 vs 10 | - | - | - | - | ns | ns | ns | ns | ns |
| | ES 0.4 vs 100 | - | - | - | - | ns | ns | ns | ns | ns |
| | ES 10 vs 100 | - | - | - | - | ns | ns | ns | ns | ns |
| | TLS | | | | | ns | ns | ns | ns | ns |

| | | | | | |
|---|---|---|---|---|---|
| 0.4 vs 10 | | | | | |
| TLS | ns | * | ns | ns | ns |
| 0.4 vs 100 | | (p=0,0130 ) | | | |
| TLS | ns | ns | ns | ns | ns |
| 10 vs 100 | | | | | |
| PTLS | ns | ns | ns | ns | ns |
| 0.4 vs 10 | | | | | |
| PTLS | ns | ns | ns | ns | ns |
| 0.4 vs 100 | | | | | |
| PTLS | ns | ns | ns | ns | ns |
| 10 vs 100 | | | | | |
| | ns | ns | ns | ns | ns |

**Table S7.** The significance of differences between the different categories of bundles in terms of apparent and net inflection point properties assessed with a one-way ANOVA followed by a Tukey post hoc (ns p>0.05, *p≤0.05, **p≤0.01, ***p≤0.001, ****p≤0.0001).

| | Strain-rate [% $s^{-1}$] | Sample | IFF [N] | IFε [%] | $IF_{pp}$ [MPa] |
|---|---|---|---|---|---|
| **App** | 0.4 | FS vs. ES | ns | ns | ns |
| | | FS vs. TLS | ns | ns | ns |
| | | FS vs. PTLS | ns | **** (p<0,0001) | ns |
| | | ES vs. TLS | ns | ns | ns |
| | | ES vs. PTLS | ns | **** (p<0,0001) | ns |
| | | TLS vs. PTLS | ns | **** (p<0,0001) | ns |
| **Net** | 0.4 | FS vs. ES | - | - | ns |
| | | FS vs. TLS | - | - | * (p=0,0498) |
| | | FS vs. PTLS | - | - | ns |
| | | ES vs. TLS | - | - | * (p=0,0253) |
| | | ES vs. PTLS | - | - | ns |
| | | TLS vs. PTLS | - | - | ns |
| **App** | 10 | FS vs. ES | ns | ns | ns |
| | | FS vs. TLS | ns | ns | ns |
| | | FS vs. PTLS | ns | ns | ns |
| | | ES vs. TLS | ns | **** (p<0,0001) | * (p=0,0491) |
| | | ES vs. PTLS | ns | **** (p<0,0001) | ns |
| | | TLS vs. PTLS | ns | **** (p<0,0001) | ns |
| **Net** | 10 | FS vs. ES | - | - | ns |
| | | FS vs. TLS | - | - | ns |
| | | FS vs. PTLS | - | - | ns |
| | | ES vs. TLS | - | - | ns |
| | | ES vs. PTLS | - | - | ns |
| | | TLS vs. PTLS | - | - | ns |

| | | | | | |
|---|---|---|---|---|---|
| **App** | 100 | FS vs. ES | ** (p=0,0010) | ns | ns |
| | | FS vs. TLS | ns | ns | ** (p=0,0026) |
| | | FS vs. PTLS | *** (p=0,0005) | ns | ns |
| | | ES vs. TLS | ns | **** (p<0,0001) | **** (p<0,0001) |
| | | ES vs. PTLS | ns | **** (p<0,0001) | ns |
| | | TLS vs. PTLS | ns | **** (p<0,0001) | **** (p=0,0216) |
| **Net** | 100 | FS vs. ES | - | - | ns |
| | | FS vs. TLS | - | - | ns |
| | | FS vs. PTLS | - | - | ns |
| | | ES vs. TLS | - | - | ** (p=0,0027) |
| | | ES vs. PTLS | - | - | ns |
| | | TLS vs. PTLS | - | - | ** (p=0,0045) |

**Table S8.** The significance of differences of the inflection points properties of bundles at different strain-rates assessed with a one-way ANOVA followed by a Tukey post hoc (ns p>0.05, *p≤0.05, **p≤0.01, ***p≤0.001, ****p≤0.0001).

| | Strain-rate [% $s^{-1}$] | IFF [N] | IFε [%] | IFσ [MPa] |
|---|---|---|---|---|
| **App** | FS 0.4 vs 10 | ns | ns | ns |
| | FS 0.4 vs 100 | **** (p<0,0001) | ns | ns |
| | FS 10 vs 100 | * (p=0,0107) | ns | ns |
| | ES 0.4 vs 10 | ns | ns | ns |
| | ES 0.4 vs 100 | ns | ns | ns |
| | ES 10 vs 100 | ns | ns | ns |
| | TLS 0.4 vs 10 | ns | ns | ns |
| | TLS 0.4 vs 100 | ns | ns | ns |
| | TLS 10 vs 100 | ns | ns | ns |
| | PTLS 0.4 vs 10 | ns | ns | ns |
| | PTLS 0.4 vs 100 | ns | ns | ns |
| | PTLS 10 vs 100 | ns | ns | ns |
| **Net** | FS 0.4 vs 10 | - | - | ns |
| | FS 0.4 vs 100 | - | - | ns |
| | FS 10 vs 100 | - | - | ns |
| | ES 0.4 vs 10 | - | - | ns |
| | ES 0.4 vs 100 | - | - | ns |

| | | | |
|---|---|---|---|
| ES<br>10 vs 100 | - | - | ns |
| TLS<br>0.4 vs 10 | | | ns |
| TLS<br>0.4 vs 100 | | | ns |
| TLS<br>10 vs 100 | | | ns |
| PTLS<br>0.4 vs 10 | | | ns |
| PTLS<br>0.4 vs 100 | | | ns |
| PTLS<br>10 vs 100 | | | ns |
| | | | ns |

**Table S9.** The significance of differences between the different categories of bundles in terms of frequency-dependent properties of scaffolds across tested strain-rates assessed with a one-way ANOVA followed by a Tukey post hoc (ns p>0.05, *p≤0.05, **p≤0.01, ***p≤0.001, ****p≤0.0001).

| Strain-rate [% $s^{-1}$] | Sample | f [Hz] | A [MPa] | λ [m] | v [m $s^{-1}$] |
|---|---|---|---|---|---|
| 0.4 | FS vs. ES | ns | ns | ns | ns |
| | FS vs. TLS | ns | *** (p=0,0004) | ns | * (p=0,0102) |
| | FS vs. PTLS | ns | ns | **** (p<0,0001) | **** (p<0,0001) |
| | ES vs. TLS | ns | * (p=0,0232) | ns | *** (p=0,0004) |
| | ES vs. PTLS | ns | ns | **** (p<0,0001) | **** (p<0,0001) |
| | TLS vs. PTLS | ns | ** (p<0,0035) | **** (p<0,0001) | **** (p<0,0001) |
| 10 | FS vs. ES | ns | ns | ns | ns |
| | FS vs. TLS | ns | **** (p<0,0001) | ns | ns |
| | FS vs. PTLS | ns | ns | ns | **** (p<0,0001) |
| | ES vs. TLS | ns | **** (p<0,0001) | ns | **** (p<0,0001) |
| | ES vs. PTLS | ns | ns | ns | ns |
| | TLS vs. PTLS | ns | **** (p<0,0001) | ns | **** (p<0,0001) |
| 100 | FS vs. ES | ns | ns | ns | ns |
| | FS vs. TLS | ns | **** (p<0,0001) | ns | **** (p<0,0001) |
| | FS vs. PTLS | **** (p<0,0001) | ns | ns | **** (p<0,0001) |
| | ES vs. TLS | ns | **** (p<0,0001) | ns | **** (p<0,0001) |
| | ES vs. PTLS | **** (p<0,0001) | ns | ns | **** (p<0,0001) |
| | TLS vs. PTLS | **** (p<0,0001) | **** (p<0,0001) | ns | **** (p<0,0001) |

**Table S10.** The significance of differences of the frequency-dependent properties of scaffolds across tested strain-rates assessed with a one-way ANOVA followed by a Tukey post hoc (ns $p>0.05$, *$p\leq0.05$, **$p\leq0.01$, ***$p\leq0.001$, ****$p\leq0.0001$).

| Sample | f<br>[Hz] | A<br>[f] | λ<br>[m] | v<br>[m $s^{-1}$] |
|---|---|---|---|---|
| FS<br>0.4 vs 10 | ns | ns | ****<br>(p<0,0001) | ns |
| FS<br>0.4 vs 100 | ****<br>(p<0,0001) | ns | ****<br>(p<0,0001) | ns |
| FS<br>10 vs 100 | ns | ns | ns | ns |
| ES<br>0.4 vs 10 | ns | ns | ****<br>(p<0,0001) | ns |
| ES<br>0.4 vs 100 | ****<br>(p<0,0001) | ns | ****<br>(p<0,0001) | ns |
| ES<br>10 vs 100 | ****<br>(p<0,0001) | ns | ns | ns |
| TLS<br>0.4 vs 10 | ns | ns | ****<br>(p<0,0001) | ns |
| TLS<br>0.4 vs 100 | ns | ****<br>(p<0,0001) | ****<br>(p<0,0001) | *<br>(p=0,0306) |
| TLS<br>10 vs 100 | ns | ****<br>(p<0,0001) | ns | ns |
| PTLS<br>0.4 vs 10 | ns | ns | ns | ns |
| PTLS<br>0.4 vs 100 | ns | ns | ns | ns |
| PTLS<br>10 vs 100 | ns | ns | ns | ns |

**Table S11.** MicroCT (µCT) measures of diameter and porosity for the various samples (PTLS; FS, ES, TLS) and the various sample's VOIs (Top, Middle, Bottom averaged on Total).

| Sample | VOI | µCT diameter (µm) | µCT porosity (%) |
|---|---|---|---|
| PTLS | Top | 490±48 | 9±6 |
| | Middle | 482±33 | 4±2 |
| | Bottom | 477±48 | 9±5 |
| | Total | 492±11 | 7±2 |
| TLS | Top | 715±68 | 30±5 |
| | Middle | 695±57 | 26±6 |
| | Bottom | 643±24 | 20±8 |
| | Total | 685±37 | 25±5 |
| ES | Top | 736±91 | 24±9 |
| | Middle | 908±87 | 36±12 |
| | Bottom | 953±53 | 37±10 |
| | Total | 866±115 | 32±7 |

| | | | |
|---|---|---|---|
| FS | Top | 1010±35 | 40±6 |
| | Middle | 992±73 | 34±9 |
| | Bottom | 988±34 | 41±11 |
| | Total | 997±12 | 38±4 |
